# Disentangling incomplete lineage sorting and introgression to refine species-tree estimates for Lake Tanganyika cichlid fishes

**DOI:** 10.1101/039396

**Authors:** Britta S. Meyer, Michael Matschiner, Walter Salzburger

## Abstract

Adaptive radiation is thought to be responsible for the evolution of a great portion of the past and present diversity of life. Instances of adaptive radiation, characterized by the rapid emergence of an array of species as a consequence to their adaptation to distinct ecological niches, are important study systems in evolutionary biology. However, because of the rapid lineage formation in these groups, and the occurrence of hybridization between the participating species, it is often difficult to reconstruct the phylogenetic history of species that underwent an adaptive radiation. In this study, we present a novel approach for species-tree estimation in rapidly diversifying lineages, where introgression is known to occur, and apply it to a multimarker dataset containing up to 16 specimens per species for a set of 45 species of East African cichlid fishes (522 individuals in total), with a main focus on the cichlid species flock of Lake Tanganyika. We first identified, using age distributions of most recent common ancestors in individual gene trees, those lineages in our dataset that show strong signatures of past introgression. This led us to formulate three hypotheses of introgression between different lineages of Tanganyika cichlids: the ancestor of Boulengerochromini (or of Boulengerochromini and Bathybatini) received genomic material from the derived H-lineage; the common ancestor of Cyprichromini and Perissodini experienced, in turn, introgression from Boulengerochromini and/or Bathybatini; and the Lake Tanganyika Haplochromini and closely related riverine lineages received genetic material from Cyphotilapiini. We then applied the multispecies coalescent model to estimate the species tree of Lake Tanganyika cichlids, but excluded the lineages involved in these introgression events, as the multispecies coalescent model does not incorporate introgression. This resulted in a robust species tree, in which the Lamprologini were placed as sister lineage to the H-lineage (including the Eretmodini), and we identify a series of rapid splitting events at the base of the H-lineage. Divergence ages estimated with the multispecies coalescent model were substantially younger than age estimates based on concatenation, and agree with the geological history of the Great Lakes of East Africa. Finally, we formally tested the three hypotheses of introgression using a likelihood framework, and find strong support for introgression between some of the cichlid tribes of Lake Tanganyika.

---

Adaptive radiation, i.e. the rapid emergence of new life-forms through the extensive diversification of an organismal lineage into new or available ecological niches, is thought to be responsible for a great deal of the extant and extinct organismal diversity on our planet (Simpson 1953; Schluter 2000; Berner and Salzburger 2015). Such outbursts of life have, for fairly long time, fascinated scientists and laymen alike, and serve as important model systems and textbook examples in evolutionary biology (see e.g. Schluter 2000; Mayr 2001; Coyne and Orr 2004). At the same time — mainly because of the rapidity of lineage formation, gene flow between the emerging species, and the frequent occurrence of convergent morphs in adaptive radiations — it has proven notoriously difficult to reconstruct the progression of adaptive radiations by means of morphological analyses and, more recently, phylogenetic reconstructions based on molecular data (e.g. Rokas and Carroll 2006; Glor 2010).

The most well known examples of adaptive radiation include Darwin’s finches on the Galapagos Islands (Grant and Grant 2008), threespine stickleback fish in the northern hemisphere (Bell and Foster 1994), anole lizards on the islands of the Caribbean (Losos 2009), and cichlid fishes in the East African Great Lakes (Fryer and Iles 1972). The exceptionally diverse cichlid assemblages in Lakes Victoria, Malawi and Tanganyika represent the most species-rich extant adaptive radiations (Salzburger et al. 2014). Hundreds of closely related cichlid species have emerged in each of these lakes in the last few millions to several thousands of years only (Kocher 2004; Seehausen 2006; Salzburger 2009; Salzburger et al. 2014), rendering the formation of new cichlid species in these lakes unusually rapid (McCune and Lovejoy 1998; Coyne and Orr 2004).

With an age of 9-12 million years, Lake Tanganyika is the oldest and, because of its great depth, also the most stable lake in Africa (Cohen et al. 1993; Salzburger et al. 2014). This has strong implications on its cichlid fauna, which is the genetically, morphologically, ecologically, and behaviorally most diverse of all extant cichlid species flocks (Salzburger et al. 2014). Lake Tanganyika is also a reservoir of more ancient lineages and the likely cradle of more modern cichlid groups (Nishida 1991; Salzburger et al. 2002b, 2005), or simply *the* ‘melting pot’ of East African cichlid diversity (Weiss et al. 2015). Understanding the cichlid assemblage of Lake Tanganyika is thus key to understand cichlid evolution in the whole of East Africa.

The cichlid fauna of Lake Tanganyika, which comprises about 200 species, has originally been divided into 12 taxonomic subgroups, so called ‘tribes’ (i.e. a taxonomic rank between subfamily and genus) (Poll 1986): Bathybatini, Cyprichromini, Ectodini, Eretmodini, Haplochromini, Lamprologini, Limnochromini, Perissodini, Tilapiini, Trematocarini, Tropheini, and Tylochromini. Takahashi (2003) revised Poll’s assignment and suggested to elevate four genera, previously included in one of Poll’s original tribes, into their own tribes: Benthochromini, Boulengerochromini, Greenwoodochromini, and Cyphotilapiini. The same author further proposed to create a new tribe for ‘*Ctenochromis^1^ benthicola* and to place Trematocarini within Bathybatini. The validity of most of these tribes is backed-up by molecular data. This is, however, not the case for Greenwoodochromini, which is nested within Limnochromini (without Benthochromini) (Duftner et al. 2005; Muschick et al. 2012) and for the new tribe containing ‘*Ctenochromis’ benthicola*, as this species forms the sister species to *Cyphotilapia* and should hence be included within Cyphotilapiini (Muschick et al. 2012; Meyer et al. 2015; Weiss et al. 2015). In addition, there is no molecular support to include Trematocarini within Bathybatini (Muschick et al. 2012; Weiss et al. 2015) and to elevate *Hemibates* into its own tribe (Weiss et al. 2015). Tropheini, on the other hand, is nested within Haplochromini (e.g. Salzburger et al. 2002b, 2005; Muschick et al. 2012; Weiss et al. 2015), so that their status as an own tribe is not justified (hence, we here use ‘Haplochromini’ as synonymous to the clade combining all haplochromines including Tropheini). Finally, the genus *Oreochromis* including its single representative in Lake Tanganyika, *O. tanganicae*, has recently been removed from Tilapiini and placed in Oreochromini (Dunz and Schliewen 2013).

Over the last quarter of a century, a number of attempts have been undertaken to resolve the phylogenetic relationships between the cichlid tribes in Lake Tanganyika (e.g. Nishida 1991; Kocher et al. 1995; Salzburger et al. 2002b; Clabaut et al. 2005; Day et al. 2008; Muschick et al. 2012; Meyer et al. 2015; Weiss et al. 2015), as well as between genera and species within tribes (see e.g. Koblmüller et al. 2004, 2010; Sturmbauer et al. 2010). Especially with respect to the placement of tribes relative to each other, there is no consensus between the different studies, which can in part be explained by the different marker sets (and phylogenetic methods) that were used at a given time. In addition, there is strong evidence for introgressive hybridization and incomplete lineage sorting in the course of the cichlid adaptive radiation in Lake Tanganyika (see e.g. Salzburger et al. 2002a; Koblmüller et al. 2010; Meyer et al. 2015; Weiss et al. 2015), both of which are expected to produce phylogenetic signals that are in conflict with the true species tree.

Nishida (1991) was the first to study tribal relationships in Tanganyikan cichlids based on genetic information, in this case allozyme data. In the resultant tree topology, the only included species of the tribe Tylochromini (*Tylochromis polylepis*) — a likely secondary colonizer of Lake Tanganyika (Koch et al. 2007) — was placed as most ancestral taxon, followed by the only member of Boulengerochromini (*Boulengerochromis microlepis*), the Bathybatini, a clade comprised by Trematocarini and Tilapiini (represented by *Oreochromis tanganicae*, another secondary colonizer), Lamprologini, and a clade that combines the remaining tribes, referred to as ‘H-lineage’ (Nishida 1991). Within this H-lineage, exclusively consisting of moutbrooding species, Nishida (1991) proposed the following relationships: Limnochromini, together with Ectodini, formed the basal branch, followed by Cyprichromini, Perissodini and a clade in which Eretmodini is placed as sister group to Haplochromini. Several studies have used mitochondrial (mt) DNA markers to address the phylogenetic relationships between cichlid tribes in Lake Tanganyika (Sturmbauer and Meyer 1993; Kocher et al. 1995; Salzburger et al. 2002b; Day et al. 2008) or a combination of mtDNA and one (Clabaut et al. 2005) or two (Muschick et al. 2012) nuclear markers. All these studies agree upon a basal position of the relatively species-poor tribes Bathybatini, Boulengerochromini, Trematocarini, as well as *O. tanganicae* and *T. polylepis* (note, however, that not all of these studies included representatives of all five taxa), yet their relative positions to one another differed (see e.g. Fig. 1 in Meyer et al. 2015). All but one (Sturmbauer and Meyer 1993) of these studies placed Eretmodini outside the H-lineage, either as sister group to Lamprologini (Kocher et al. 1995; Clabaut et al. 2005; Day et al. 2008) or as sister group to a clade combining Lamprologini and the remaining taxa of the H-lineage (Salzburger et al. 2002b; Muschick et al. 2012). The relative order of taxa within the H-lineage varied between the different studies (see e.g. Fig. 1 in Meyer et al. 2015).

**Figure 1:**
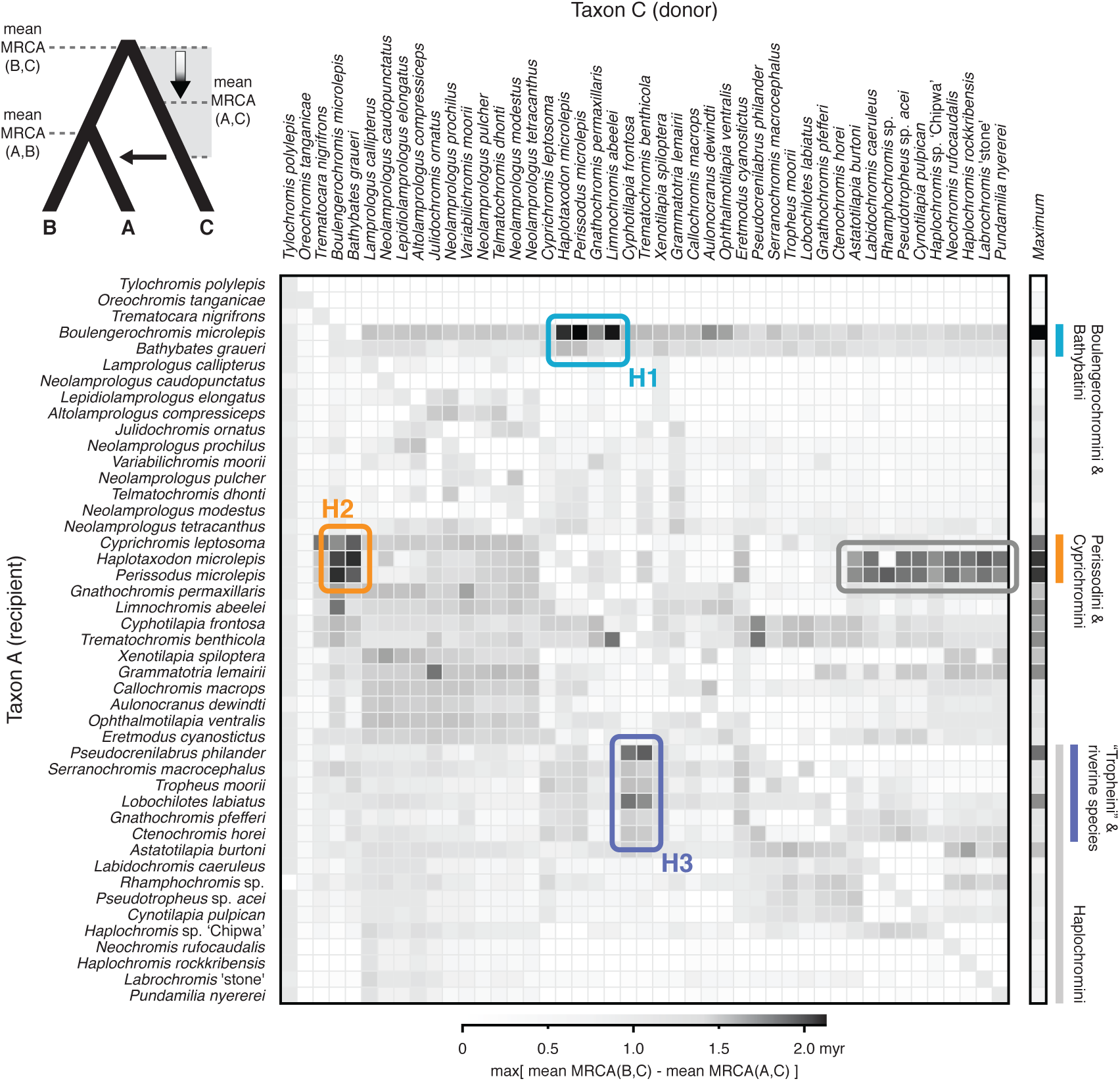
Differences in mean MRCA age estimates in three-taxon comparisons. The insert at the top left indicates how introgression from taxon C to taxon A can decrease the mean age estimate for the MRCA of A and C, compared to the mean MRCA age estimate between taxa B and C. Heat map cells indicate the maximum age difference between these mean MRCA age estimates, with taxa A and C of the three-taxon comparison according to cell row and column, and any of the remaining species as taxon B. The darkest heat map cells indicate differences in mean MRCA age estimates of up to 2.13 myr and serve as the basis for three hypotheses of introgression, H1-H3, marked by colored frames (see text). The frame in gray indicates a fourth possible introgression event between Haplochromini and Perissodini. For each species, the maximum age difference in mean MRCA age estimates, in all possible three-taxon comparisons that include this species as taxon A, is shown in the column to the right of the heat map.

More recently, the problem of the tribal relationships within the cichlid assemblage of Lake Tanganyika has been addressed using larger numbers of nuclear DNA markers. Friedman et al. (2013) used 10 nuclear markers to resolve phylogenetic relationships in a much larger context, but had most Tanganyikan cichlid tribes represented in their data set. In their phylogeny, *T. polylepis* appeared as the most ancestral cichlid lineage in Lake Tanganyika, followed by *O. tanganicae*, Bathybatini, Boulengerochromini, a clade formed by Lamprologini as sister group to the combined Cyprichromini, Perissodini, Cyphotilapiini, and Limnochromini, and a clade comprised by Ectodini as sistergroup to Eretmodini and Haplochromini. Meyer et al. (2015), using sequence information from 42 nuclear loci, presented a phylogenetic hypothesis, in which, again, *T. polylepis* and *O. tanganicae* branched off first, followed by a clade formed by Bathybatini and Trematocarini, while Boulengerochromini were placed as sister group to Lamprologini and the H-lineage. Therein, a clade comprising Perissodini and Cyprichromini was recovered as sister taxon to all other H-lineage tribes, Cyphotilapiini clustered with Limnochromini, and Eretmodini formed a clade with Haplochromini. However, when inspecting concordance between individual gene trees, certain discrepancies were found. For example, the placement of Eretmodini as sister group to Haplochromini was only supported in a subset of 14 congruent gene trees, whereas another 13 gene trees supported Eretmodini as most basal taxon within the H-lineage (Meyer et al. 2015). McGee et al. (2016), in a comparison of jaw morphology between cichlids from Lake Tanganyika and Malawi, used sequence information from a large number of ultraconserved elements. In their phylogeny containing sequence information from 56 Tanganyika and Malawi cichlids representing 10 different tribes (and using a representative of Bathybatini as outgroup), the Lamprologini were placed as sistergroup to a clade containing the H-lineage taxa, in which two sub-clades were recovered: A clade with Perissodini and Cyprichromini as sistergroup to Ectodini and Cyphotilapiini with Limnochromini, and a clade with Eretmodini and Haplochromini (McGee et al. 2016).

In contrast to the above studies, Weiss et al. (2015) used AFLP markers to resolve phylogenetic relationships between the cichlid tribes of Lake Tanganyika, and to place them into a larger phylogenetic context. Their neighbor-joining consensus phylogeny (not including *T. polylepis* and using Oreochromini as outgroup) suggests that Trematocarini form an independent lineage, separated from the remaining tribes by riverine Austrotilapiines. A clade formed by Bathybatini and Boulengerochromini is then placed as sister group to the Lamprologini and the H-lineage; followed by a clade comprised by Ectodini, Limnochromini, Cyprichromini, Benthochromini, Perissodini, and Cyphotilapiini; and the Eretmodini as sister group to the Haplochromini. The study of Weiss et al. (2015) is, thus, the first to suggest monophyly of what they call ‘ancient Tanganyika mouthbrooders’ (all members of the H-lineage except Eretmodini and Haplochromini).

Taken together, while the general phylogenetic structure of the cichlid assemblage of Lake Tanganyika is well established — with secondary colonizers of Tylochromini and Oreochromini as most ancestral splits, and Trematocarini, Bathybatini and Boulengerochromini as ancestral lineages that are sister to a clade formed by the Lamprologini and the H-lineage — several areas of uncertainty remain (see Meyer et al. 2015). For example, the relative position of Trematocarini, Bathybatini and Boulengerochromini to one another is still unclear, as is the relative position of the H-lineage taxa to one another (for example the placement of Eretmodini).

In this study, we take a novel approach to address the phylogenetic relationships between the cichlid tribes of East African Lake Tanganyika. We use 40 nuclear markers established in Meyer and Salzburger (2012) and Meyer et al. (2015). Yet, instead of using a single representative per species as in Meyer et al. (2015), we use up to 16 specimens per species, resulting in nuclear DNA sequences from 522 specimens. This allows us to first disentangle the signals of introgression and incomplete lineage sorting among cichlid taxa, using new methodology to detect gene flow from population-level and species-level data sets. We then apply the multispecies coalescent model to identify the species tree of Lake Tanganyika cichlid lineages after filtering out signals of introgression that would represent a violatation of this model. Finally, by comparing age estimates recovered with the multispecies coalescent model and with concatenation, we reconcile phylogenetic divergence-date estimates of cichlids with the geological history of their environment, to build a coherent timeline of cichlid diversification in the Great Lakes of East Africa.

## Materials and Methods

### Data collection

All specimens of cichlid fishes from Lake Tanganyika used in this study were collected in accordance with the national legislation of the Republic of Zambia and under the memorandum of understanding (MOU) between the involved institutions (University of Basel, Switzerland, University of Zambia, Lusaka, Zambia, and Department of Fisheries, Lake Tanganyika branch, Mpulungu, Zambia). Fish collections were conducted as described in Muschick et al. (2012). We used up to 16 individuals from 45 different cichlid species (a total of 522 individuals). Our sampling design comprised representatives of all major cichlid lineages of the Lake Tanganyika radiation (33 species) plus four representatives each of Lake Victoria and Lake Malawi, and four riverine taxa. The gross of samples originated from fieldtrips to Zambia, i.e. to Lake Tanganyika and the Kafue River in the years 2007, 2008 and 2011. Further samples were aquaria-bred at the Swiss Federal Institute of Aquatic Science and Technology (EAWAG; these were kindly provided by Ole Seehausen) and at the University of Basel. The number of sampled individuals per species and their place of origin are listed in Supplementary Tables S1-2. Fin clips were stored at −20 °C in 100% ethanol until further processing.

Extraction of DNA from ethanol-preserved fin clips was performed with a Qiagen Biosprint 96 robot following the manufacturer’s protocol (QIAGEN, Hombrechtikon, Switzerland). For multiplexed PCR amplifications, we used a primer set consisting of 40 markers (encompassing exons, introns, and untranslated regions of nuclear coding genes) as described in Meyer and Salzburger (2012) and Meyer et al. (2015) (see Table 1 for further details). PCR amplifications and library preparations (including several cleaning and pooling steps) were carried out as described in Meyer et al. (2015); pyrosequencing was performed on the GS FLX system (454 Sequencing, Roche). The sequencing design in the current study differs from that of Meyer et al. (2015), who included only a single specimen for each species (instead of up to 16 as in the current study) and used a consensus sequence (instead of the two alleles as in the current study).

**Table 1:**
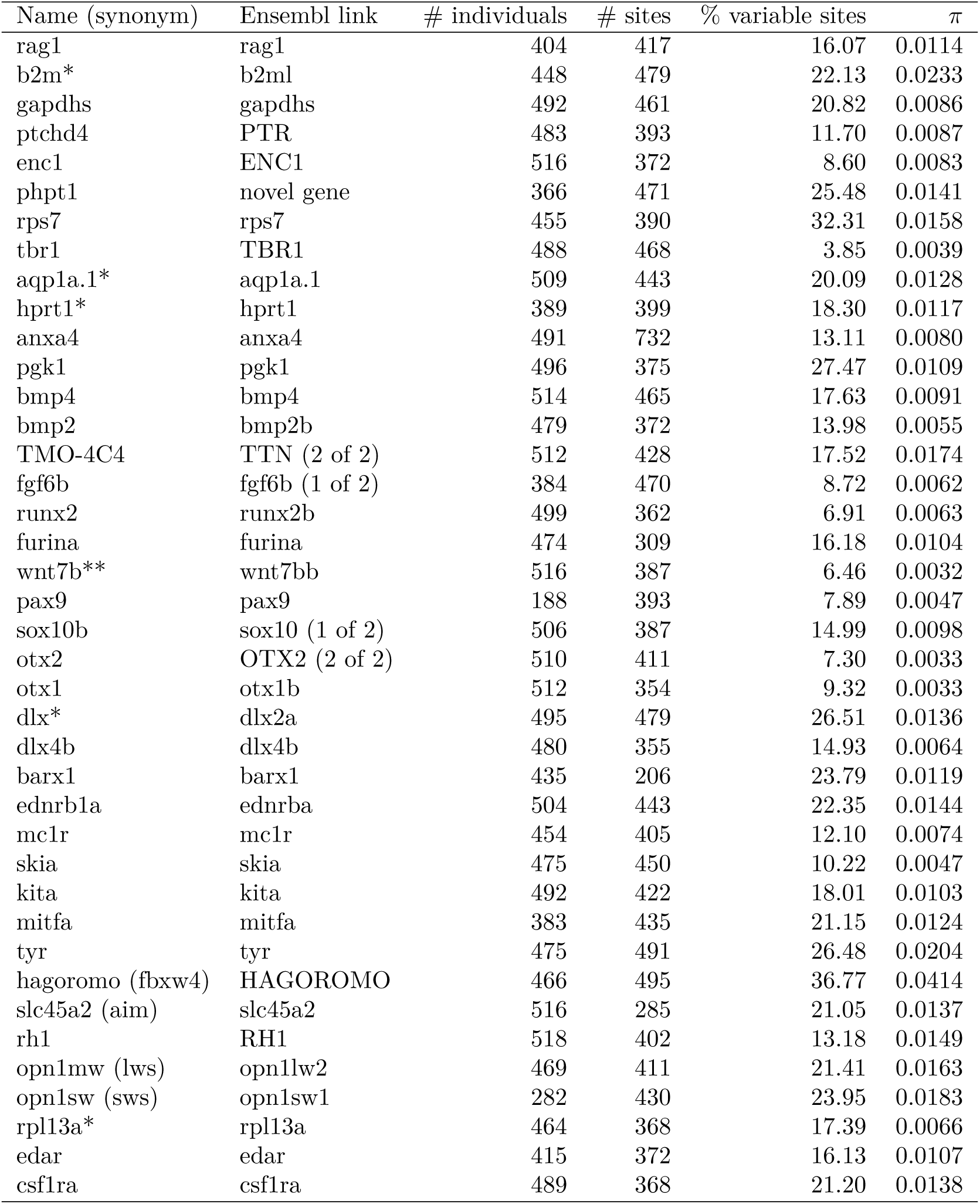
Description of the phylogenetic markers used in this study. The marker name, the link to the corresponding Ensembl entry for Tilapia, the number of sequenced individuals, the alignment length of each marker, the proportion of variable sites, and the genetic diversity (n) of each marker are given. *Marker excluded from all gene-tree analyses and MRCA age comparisons due to missing data in *Tylochromis polylepis.* **Unconstrained gene-tree analyses failed for this marker, and it was thus excluded from MRCA age comparisons.

The generated sequence reads were demultiplexed and assembled using Roche’s sffinfo tool, PRINSEQ (Schmieder and Edwards 2011), bwa (Li and Durbin 2010), and Geneious (Biomatters Ltd, Auckland, New Zealand; available from www.geneious.com) with the same versions and settings as in Meyer et al. (2015). Due to the long read length provided by the 454 technology, phased alleles could be determined in all individuals. Thus, we obtained between 376 and 1036 (mean 918.9) sequences per marker (Table 1). Alignments for each marker containing all alleles from all individuals were obtained with the software MAFFT v.7.017 (Katoh and Standley 2013), using the “-auto” option to automatically detect the appropriate alignment strategy. The alignments were inspected visually and were locally corrected where necessary (Supplementary File S1).

### Gene-tree inference

As our phylogenetic data set included both intra-and interspecific sequence variation, we performed phylogenetic inference using the multispecies coalescent model of *BEAST (Heled and Drummond 2010), implemented in the Bayesian phylogenetic inference software BEAST v.2.1.3 (Bouckaert et al. 2014). However, we expected that gene tree topologies could be discordant not only due to incomplete lineage sorting (ILS), but potentially also due to introgression. As the *BEAST model only accounts for ILS but excludes introgression, we conducted separate *BEAST analyses for each marker, assuming that recombination only takes place between markers, but not within. Phylogenetic inference was performed using an HKY model (Hasegawa et al. 1985) of sequence evolution and the Yule process (Yule 1925) as the species-tree prior. For computational reasons, we assumed no among-site rate heterogeneity and used empirical base frequencies. To time calibrate the phylogeny, we used a strict molecular clock model and two fossil-based node age prior distributions. Based on the time-calibrated phylogeny of cichlid fishes by McMahan et al. (2013), we constrained the divergence of Tylochromini and Austrotilapiini between 54.8 and 33.0 million years ago (Ma) (95% confidence interval; mean = 42.1 Ma), and that of Oreochromini and Austrotilapiini between 31.6 and 17.3 Ma (mean = 23.3 Ma). In our data set, Tylochromini were represented by *Tylochromis polylepis*, Oreochromini were represented by *Oreochromis tanganicae*, and 43 other included species were members of Austrotilapiini, which were assumed to be monophyletic. Both time constraints were implemented as log-normal prior distributions, with offsets 18.2 and 8.2 Ma, means (in real space) 23.894 and 15.067 Ma, and standard deviations 0.231 and 0.240 Ma, respectively. For consistency between ages estimated with different markers, we excluded five out of 40 markers due to missing sequence data for *Tylochromis polylepis* (see Table 1). We conducted five replicate analyses for each of the 35 markers, with 4 billion Monte Carlo Markov chain (MCMC) generations of which the first ten percent were discarded as burn-in. Run convergence was assessed by comparison of traces within and between run replicates and by effective sample sizes (ESS) greater than 100 for each model parameter. For each marker, 100 time-calibrated gene trees were sampled from the joint posterior distribution of the five replicate analyses. Each of these gene trees was further pruned to include only a single randomly chosen individual per species. This was repeated 10 times per tree to produce posterior sets of 1000 species-level gene trees for each marker.

### Detection of putatively introgressed lineages

*Comparison o mean MRCA age estimates.* — Posterior samples of species-level gene tree distributions were used to calculate mean age estimates of most recent common ancestors (MRCA) in all pairwise species comparisons. As gene-tree inference for one out of 35 markers (wnt7b) failed to converge, we used gene tree distributions of 34 markers for these calculations. Following Marcussen et al. (2014), we expected that introgression could be detectable through MRCA age estimates in three-taxon comparisons (see Fig. 1 for illustration): for a given set of taxa A, B, C, two of the three mean MRCA age estimates should be similar if introgression is absent. For example, if A and B are sister lineages and C is the outgroup, then the MRCA for A and C and the MRCA for B and C should have a similar age, even in the presence of ILS (assuming comparable population sizes and mutation rates for A and B). However, if part of the genome of A is affected by introgression coming from C, then the mean MRCA age for A and C should be younger than the MRCA for B and C. Thus, to identify the species with the strongest signals of introgression, we compared mean MRCA age estimates in each possible three-taxon combination, testing each of the positions A, B, and C for each of the 45 species in our data set (a total of 45^3^ comparisons). We recorded the difference between the age estimate of the MRCA of B and C and the age estimate of the MRCA of A and C if (i) the MRCA of A and C appeared younger than that of B and C, and if (ii) the MRCA of A and B was the youngest of the three, indicating that A and B are sister groups in this three-taxon comparison. If these conditions were not met, the age difference was recorded as zero. The results of these three-taxon comparisons were summarized by calculating, for each pair of the 45 species, the maximal age difference found with these two species in positions A and C, and with any other species in position B. For each of the 45 species, we also calculated the maximal age difference when this species was in position A, and any other two species were in positions B and C (thus the maximal age difference in 45^2^ comparisons).

Based on the maximal age differences per species pair and per individual species, we developed three hypotheses of introgression between Lake Tanganyika cichlid lineages (Fig. 1; Supplementary Table S3) that were to be tested in subsequent analyses: H1; an early representative of Boulengerochromini, or a common ancestor of Boulengerochromini and Bathybatini, received, through introgression, genomic material from members of the H-lineage, most likely from the Perissodini or Limnochromini; H2; an early member or common ancestor of the Perissodini and Cyprichromini experienced introgression from the Boulengerochromini or Bathybatini; H3; a common ancestor of the Lake Tanganyika Haplochromini (i.e. the “Tropheini”) plus the riverine lineages *Pseudocrenilabrus* and *Serranochromis* experienced introgression from an ancestor or early member of Cyphotilapiini. A fourth potential introgression event was indicated by high maximal age differences in comparisons involving Haplochromini of lakes Malawi and Victoria (or one of *Astatotilapia burtoni* and *Haplochromis* sp. ‘Chipwa’) as taxon C and Perissodini as taxon A (see Fig. 1). However, as these signals for introgression between the Haplochromini and Perissodini were not as strong as those for introgression between the Boulengerochromini or Bathybatini and Perissodini (H2), we did not further investigate introgression between the Haplochromini and the Perissodini, and instead focussed on testing the first three hypotheses H1-H3.

*f-statistics*.— The *f*_4_-statistic, introduced by Reich et al. (2009), is a powerful measure to distinguish introgression from incomplete lineage sorting, based on allele frequencies of four populations. With populations A, B, C, and D, and the assumed unrooted population topology (A,B),(C,D), the *f*_4_-statistic is calculated as the product of the difference of allele frequencies between A and B, and between C and D. While this statistic was initially applied to populations of a single species or sister species (Reich et al. 2009), it has been shown to be effective also to detect introgression between species that have diverged several millions of years ago (Martin et al. 2015). We thus calculated the *f*4-statistic for sets of four taxa according to our hypotheses of introgression outlined above. In order to test for introgression between the H-lineage and the Boulengerochromini or Bathybatini (hypothesis H1), we assumed an unrooted topology (A,B),(C,D), where A was one of the two taxa *Oreochromis tanganicae* and *Trematocara nigrifrons*, B was *Boulengerochromis microlepis* or *Bathybates graueri*, C was either *Perissodus microlepis* or *Limnochromis abeelei*, and D was a member of Lamprologini, either *Lepidiolamprologus elongatus* or *Neolamprologus prochilus.* In these comparisons, the two members of Lamprologini were chosen as this tribe, which is not included in the H-lineage, showed no or very weak signals of introgression into the Boulengerochromini and Bathybatini based on MRCA age estimates, in contrast to the Limnochromini and Perissodini. To test for introgression between the Boulengerochromini or Bathybatini and the Perissodini and Cyprichromini (H2), we repeated this test, again with either *Oreochromis tanganicae* or *Trematocara nigrifrons* as taxon A, *Boulengerochromis microlepis* or *Bathybates graueri* as taxon B, and now *Perissodus microlepis* or *Cyprichromis leptosoma* as taxon C, and *Grammatotria lemairii* or *Ophthalmotilapia ventralis* as taxon D. Finally, introgression between Cyphotilapiini and a clade combining Lake Tanganyika Haplochromini with riverine lineages (H3) was assessed with one of the two members of Ectodini as species A, a member of Cyphotilapiini as species B, either *Pseudocrenilabrus philander* or *Lobochilotes labiatus* as species C, and *Astatotilapia burtoni* or *Haplochromis* sp. Chipwa’ as species D.

In the absence of introgression, the *f*4-statistic is expected to be zero, regardless of whether ILS is present or not. Thus, introgression between one of the two species A and B and one of the species C and D can be inferred if the *f*_4_-statistic is significantly different from zero (Reich et al. 2009). Whether or not this is the case is usually assessed on the basis of standard errors calculated through a block jackknife procedure. The use of jackknife standard errors for confidence interval estimation assumes that the underlying data is normally distributed, however, this may often not be the case for the *f*4-statistic, especially with more divergent species-level allele frequency data. This is because with more divergent populations, a larger numbers of single nucleotide polymorphisms (SNPs) will be fixed within populations but different between them. As a result, the *f*_4_-statistic will be exactly zero for a large number of SNPs. If jackknife blocks include a large number of linked sites, the per-block *f*4-statistic may then also be close to zero more often than assumed under normality.

To use the *f*4-statistic as a test of introgression with our population-level multi-marker data set, we therefore applied not only a block jackknife procedure but also developed a new approach to assess significance, which does not assume normality and accounts for linkage of genetic variation within markers. To this end, we conducted simulations to evaluate how often the observed *f*_4_-statistic can be reproduced in the absence of introgression, based on ILS alone. We used the coalescent software fastsimcoal2 v.2.5.2 (Excoffier et al. 2013) to produce sequence data sets for four populations that are similar to the true data set in terms of size and amount of missing data. Simulations were carried out using a wrapper script for fastsimcoal2, and simulation parameters for effective population sizes and divergence times were optimized during a burn-in phase. The burn-in phase was stopped as soon as parameter combinations were found with which the resulting simulated sequence variation matched the observed sequence variation across all markers in the proportion of SNPs that are variable in more than one species and in the proportion of SNPs that are variable within both pairs of species. Subsequent to burn-in, 1000 sets of coalescent simulations were carried out, where in each of these sets sequence data was simulated separately for all 39-40 markers included in a given four species comparison. For each comparison, we used only sites that were bi-allelic among the four species included in the comparison. We interpreted the observed *f*4-statistic as evidence for introgression if less than 5% of the 1000 data sets simulated without introgression produced *f*4 values at least as extreme as the observed. Our simulation-based test procedure was implemented in the new software F4, which is available at https://github.com/mmatschiner/F4.

### Species-tree inference with reduced taxon sets

As some of the species showed stronger signals of introgression than others, we assumed that excluding the species with the strongest signals would produce a largely introgression-free reduced taxon set, for which the multispecies coalescent model of *BEAST should be appropriate in a joint analysis of all markers. Based on signals of introgression observed with gene tree MRCA age comparisons and the *f*4-statistic, we excluded the following taxa from species-tree inference with *BEAST: *Boulengerochromis microlepis, Bathybates graueri*, members of Perissodini and Cyprichromini, the riverine lineages *Pseudocrenilabrus* and *Serranochromis*, as well as all Lake Tanganyika haplochromines except *Astatotilapia burtoni* and *Haplochromis* sp. ‘Chipwa’ (i.e. only the “Tropheini”). After excluding all individuals of these species, our data set contained population-level data for 34 remaining species. As we did for gene-tree analyses with *BEAST, we again used the Yule process as the species-tree prior and an HKY model of sequence evolution without among-site rate variation and base frequencies as empirically observed. Time calibration was again based on a strict molecular clock model with marker-specific clock rates, and on the same two time constraints for the ages of Tylochromini and Oreochromini. Per-branch effective population sizes were estimated as part of the analyses, however, for computational reasons, the data set was reduced to include maximally four randomly selected phased sequences per species for each marker. To assess robustness of results, random sequence selection was performed twice, and the two resulting data sets were used for independent sets of *BEAST analyses. For each data set, we performed four replicate *BEAST runs with 4 billion MCMC steps per replicate, of which the first 40% were discarded as burn-in.

To test how the application of the multispecies coalescent model influences the estimated species-tree topology and timeline of cichlid diversification, we also performed BEAST analyses with concatenated marker sets for the 34 putatively introgression-free species. It has been argued that concatenation is a special case of the multispecies coalescent model in which all gene trees are constrained to be identical in topology and branch lengths (Edwards et al. 2016). Thus, by comparing results based on concatenation and the multispecies coalescent model, we effectively assess how the release of the constraint of identical gene trees for all markers affects the species-tree estimate. We performed Bayesian analyses of concatenated marker sets using the same model of sequence evolution, molecular clock, and speciation, as for species-tree inference with the multispecies coalescent model of *BEAST. For analyses of concatenated marker sets, we removed within-species sequence variation by random selection of a single sequence per marker and species. This step was repeated twice to assess the effect of stochasticity in the random sequence selection. For the two generated data sets, we conducted five replicate BEAST analyses, each using 500 million MCMC generations, of which we discarded the first 20% as burn-in.

Run convergence in all BEAST analyses was evaluated by ESS values above 100 for all model parameters, and by visual inspection of parameter traces within and between replicates. Posterior tree samples of run replicates were combined to produce maximum clade credibility (MCC) species trees with the software TreeAnnotator v.2.1.3 (Bouckaert et al. 2014), with node heights set to mean age estimates.

### Constrained gene-tree inference based on the species tree

We now used the species tree based on the putatively introgression-free reduced taxon set to obtain more reliable estimates of gene trees, which were to be used for likelihood inference of introgression (see below). These new gene-tree analyses were performed separately for each of 35 markers, again excluding five markers for which no sequence data for *Tylochromis polylepis* were available. Constrained gene trees were inferred with a new set of *BEAST analyses, with additional constraints based on the previously inferred species tree. These constraints were placed on the age and topology of 15 clades that were supported with Bayesian posterior probabilities (BPP) of at least 0.99 in the species tree. To constrain the age of these clades according to the previously inferred species tree, we used lognormal prior distributions fitted to the distribution of posterior age estimates from the previous analysis. However, the topology of these clades could not simply be constrained as monophyletic with respect to all other taxa, as the new set of gene-tree analyses included additional species (see below) that might fall into these clades. To allow inclusion of additional taxa within these clades, we used “CladeConstraint” priors from the Sampled Ancestors (Gavryushkina et al. 2014) package for BEAST. These priors allow specifications of ingroup and outgroup taxa for a clade, and all taxa not listed in either of the two categories are allowed to be included in one or the other. Data sets used for these analyses included maximally four sequences per species, for the same set of 34 species that were included in the inference of the putatively introgression-free reduced species tree, and for one of the three species *Boulengerochromis microlepis, Perissodus microlepis*, and *Pseudocrenilabrus philander.* Three sets of analyses were conducted so that each included only one of these three species, and within-species sequence data was again selected at random, which was repeated twice to produce two equivalent data sets for separate analyses. Thus, for each of the 35 markers, we used six data sets that differed slightly in species sets and selected sequences per species. Per marker, three replicate *BEAST analyses were carried out for each of the six data sets, with the same settings as for the previous species-tree analyses, except that only 500 million MCMC generations were used for each analysis replicate. After run convergence was assessed, we merged the posterior tree distributions for each set of three run replicates. For each marker, and for each of the data sets used for this marker, we produced summary trees with only one tip per species by randomly removing all but one individual per species from each tree of the posterior distribution before generating MCC gene trees. Since visual inspection showed that for each marker and species set, the two MCC gene trees resulting from repeated random selection of within-species sequence data were highly congruent, we generated joint MCC gene trees from the combined posterior tree distributions of the two analyses.

### Likelihood-based tests for introgression

We used the inferred sets of constrained gene trees to test the three hypotheses of introgression, H1-H3, in a likelihood framework. For the three sets of taxa that each included one of the three species with strong signals of introgression (*Boulengerochromis microlepis, Perissodus microlepis*, and *Pseudocrenilabrus philander*), the inferred MCC gene trees were used jointly to assess the phylogenetic position and the most probable source of introgression into this species. We used function “InferNetwork_ML” (Yu et al. 2014) of the program PhyloNet v.3.5.6 (Than et al. 2008) to compare the likelihoods of the set of gene trees under ILS alone, and after adding a single introgression edge to the species tree. For computational reasons, and because we expected introgression into *Boulengerochromis microlepis, Perissodus microlepis*, and *Pseudocrenilabrus philander* to originate from basal branches or stem lineages of tribes rather than from their crowns, we reduced all gene trees to single representatives of tribes for these analyses. As representatives of tribes, of which our taxon set included multiple species, we chose the lamprologine *Neolamprologus prochilus*, the ectodine *Ophthalmotilapia ventralis*, the limnochromine *Gnathochromis permaxillaris*, the cyphotilapiine *Trematochromis benthicola*, and the haplochromine *Pundamilia nyererei*, due to low amounts of missing data for these taxa and their nested positions in the respective tribes. After further removing two of the 35 gene trees due to missing data for Trematocarini, all 33 remaining gene trees included a tip for each of ten tribes. To disregard topologically uncertain nodes in gene trees, we applied a gene trees support threshold of BPP ≥ 0.9 in all PhyloNet analyses (option “-b”). Ten replicate analyses were performed for all sets of gene trees (option “-x”) to assess convergence, and branch lengths and inheritance probabilities of introgression edges were optimized after the topology search (option “-po”).

## Results

### Detection of putatively introgressed lineages

*Comparison of mean MRCA age estimates*.— Our comparison of MRCA mean age estimates in 35 gene trees showed that divergences in three-taxon comparisons were the least tree-like when members of three groups of Lake Tanganyika cichlids were included. These groups comprised (i) the Boulengerochromini and Bathybatini, (ii) the Perissodini and Cyprichromini, and (iii) Lake Tanganyika Haplochromini and the riverine lineages *Pseudocrenilabrus* and *Serranochromis* (Fig. 1). When *Boulengerochromis microlepis* was included in the comparison as taxon A (see scheme in Fig. 1), the MRCA age estimates between taxa A and C were up to 2.13 myr younger those between taxa A and B, which was the case in a comparison with *Trematocara nigrifrons* as taxon B and *Perissodus microlepis* as taxon C (Supplementary Table S3). For the three species involved in this comparison, *Boulengerochromis microlepis* was closest to *Trematocara nigrifrons* in the MCC gene trees of 15 markers, closest to *Perissodus microlepis* in the MCC gene trees of 12 markers, and ancestral to the other two taxa in the MCC gene trees of 5 markers (the trees of the remaining markers trees did not include one of these species) (Supplementary File S2). Comparable differences in MRCA age estimates were observed when other members of the Perissodini or Limnochromini were included in the comparison instead of *Perissodus microlepis* (Fig. 1), or when *Bathybates graueri* was used instead of *Boulengerochromis microlepis.*

When members of Cyprichromini and Perissodini were used as taxon A in three-taxon comparisons, the greatest difference between the mean age estimate for the MRCA of taxa B and C and the mean age estimate for the MRCA of taxa A and C was found in the species trio *Perissodus microlepis* (as taxon A), *Gnathochromis pfefferi* (as taxon B), and *Boulengerochromis microlepis* (as taxon C). The mean MRCA age estimate of *Perissodus microlepis* and each of the other two species was about 2 myr younger than that between them (Supplementary Table S3). *Perissodus microlepis* was found closer to *Gnathochromis pfefferi* in the MCC gene trees of 18 markers, and closer to *Boulengerochromis microlepis* in the MCC gene trees of 15 markers, and it was the outgroup to the other two taxa in the remaining 5 MCC gene trees (Supplementary File S2). Very similar patterns were found when other members of Perissodini and Cyprichromini were used instead of *Perissodus microlepis*, or when *Boulengerochromis microlepis* was replaced by *Bathybates graueri* (Fig. 1).

We further observed large differences between the mean age estimate for the MRCA of taxa B and C and the mean age estimate for the MRCA of taxa A and C in comparisons involving Lake Tanganyika haplochromines, *Pseudocrenilabrus*, or *Serranochromis* as taxon A, together with a representative of the Cyphotilapiini as taxon C (Fig. 1). These differences were most pronounced in a comparison including *Pseudocrenilabrus philander* as taxon A, *Neochromis rufocaudalis* as taxon B, and *Trematochromis benthicola* as taxon C, where the mean MRCA age estimate for *Trematochromis benthicola*, and *Neochromis rufocaudalis* was about 1.8 myr older than both mean age estimates for MRCA involving *Pseudocrenilabrus philander* (Supplementary Table S3). Among these three species, *Pseudocrenilabrus philander* was sister to *Neochromis rufocaudalis* in the MCC gene trees of 15 markers and to *Trematochromis benthicola* in the MCC gene trees of 12 markers, and *Neochromis rufocaudalis* and *Trematochromis benthicola* were most closely related in the remaining 7 MCC gene trees (Supplementary File S2).

Taken together, these results are consistent with at least three hypothesized introgression events during the early diversification of Lake Tanganyika cichlids: MRCA age differences and gene topology frequencies in three-taxon comparisons involving *Boulengerochromis microlepis* might be explained by introgression from the H-lineage to the Boulengerochromini or to a common ancestor of Boulengerochromini and Bathybatini (hypothesis H1), introgression may further have occurred between a member of the Boulengerochromini and Bathybatini, and an ancestor of Perissodini and Cyprichromini (hypothesis H2), and *Pseudocrenilabrus* or an early representative of Lake Tanganyika haplochromines may have experienced introgression from the Cyphotilapiini (hypothesis H3).

*f-statistics*.— The *f*_4_-statistic was calculated for a total of 46 four-taxon comparisons, with between 390 and 745 bi-allelic SNPs that could be extracted from our multi marker data set (Supplementary Table S4). Between 2.0 and 5.4% of these SNPs were variable within both pairs of sister taxa, indicating gene flow between lineages that are separated by millions of years. Most of the calculated *f_4_* values (39 out of 46) were negative, suggesting gene flow according to the three tested hypotheses of introgression, H1-H3. Based on a block jackknife procedure with a block size of 20 SNPs, *f*_4_ values were significantly different from zero in six comparisons; however, the assumption of normality of jackknife block *f_4_* values was violated in all of them (Shapiro-Wilks test for normality, *p* ≤ 0.02). This indicates that a block jackknife procedure may not be suitable to assess significance of the *f*4-statistic. By simulating data sets without introgression that were comparable to the observed data set in terms of SNP number and variation, we found that per four-taxon comparisons, between 11% and 49% of these data sets produced *f_4_* values as extreme or more extreme than the empirically observed *f*4 value (see Supplementary Table S4).

### Species-tree inference with reduced taxon sets

Recall, that in order to test for robustness of phylogenetic inference, we performed all species-tree analyses with two different data sets resulting from random selection of sequences as representatives of within-species variation. For each set of analyses, these two data sets always produced two near-identical phylogenetic estimates of the interrelationships between Lake Tanganyika cichlid tribes. Rather than discussing these two phylogenies separately, we thus combined the two posterior tree samples for each of these analyses to produce joint MCC phylogenies as our best species-tree estimates.

Species-tree inference for the 34 taxa with no or only weak signs of introgression produced a strongly supported topology (mean BPP = 0.83) (Fig. 2; Supplementary File S2). In this species tree, the Trematocarini are recovered as the sister group to all other Tanganyikan cichlid tribes (BPP = 1.0), with the exception of the Tylochromini and Oreochromini that invaded the lake secondarily (Salzburger et al. 2002b). We found strong support (BPP = 1.0) for reciprocal monophyly of the Lamprologini and a clade combining all members of the H-lineage. The monophyly of each tribe is equally well supported, as is the sister-group relationship between Limnochromini and Cyphotilapiini, and the monophyly of both the Lake Victoria and Lake Malawi cichlids (BPP = 1.0). Overall, the tree topology corroborates the relationships recently identified with a concatenated analysis of a similar marker set (Meyer et al. 2015), but is unable to resolve the sequence of divergences between the Ectodini, the combined Cyphotilapiini and Limnochromini, the Eretmodini and the Haplochromini. Bayesian time calibration indicates that these four groups have diverged near-instantaneously around 7 Ma (95% highest posterior density intervals (HPD) 9.1-5.3 Ma) (Fig. 2).

**Figure 2:**
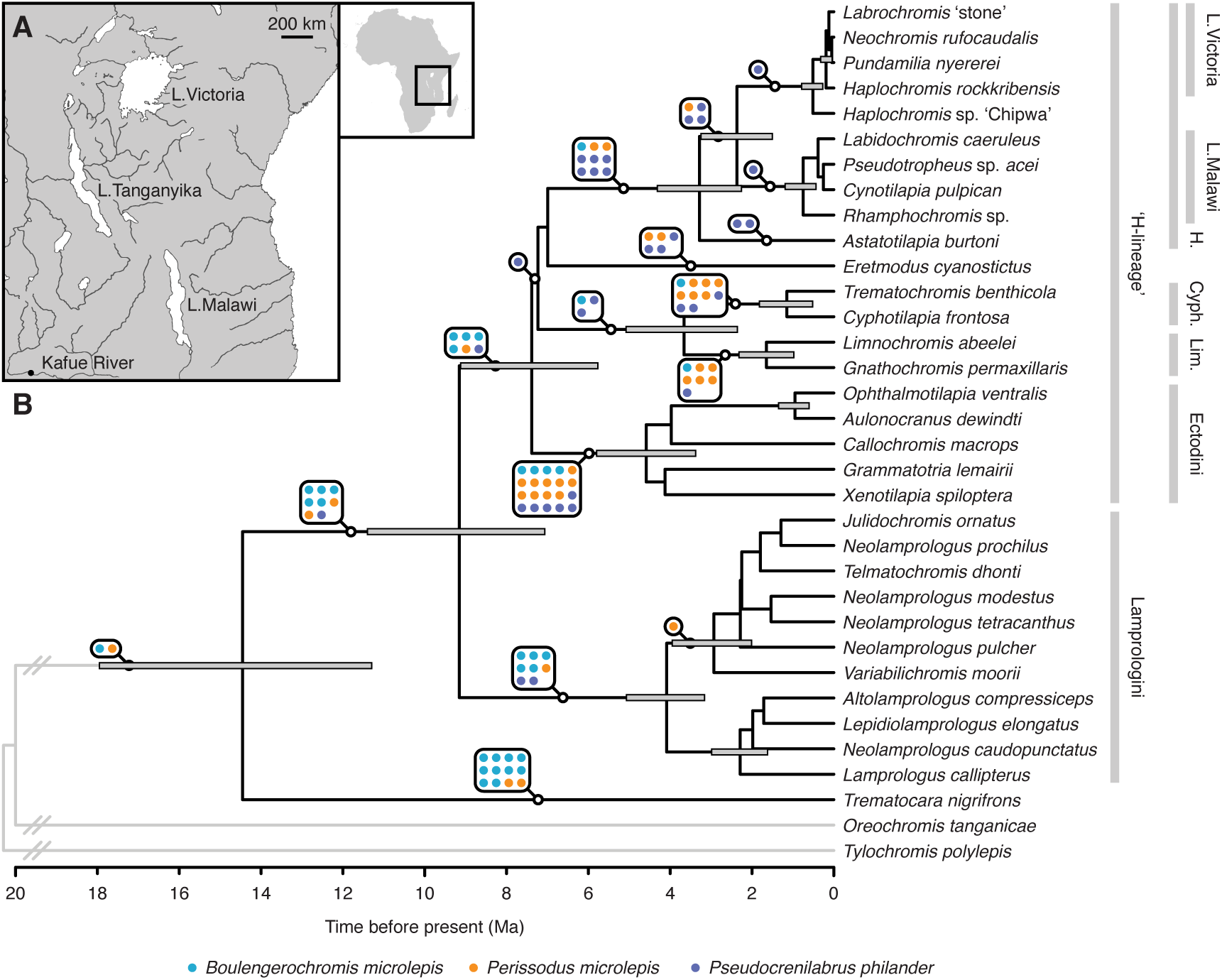
Species tree for East African cichlid lineages. A) Map of East Africa showing the three great lakes Tanganyika, Malawi, and Victoria. B) Time-calibrated species tree for a reduced taxon set of 34 East African cichlid species with no or weak signals of introgression, based on 35 markers and inferred using the multispecies coalescent model. Node bars indicate 95% HPD intervals for clades supported with a BPP of 0.99 or higher. Turquoise, orange, and blue dots on branches indicate how often *Boulengerochromis microlepis, Perissodus microlepis* and *Pseudocrenilabrus philander*, respectively, were found to attach to these branches (or to descendent branches) in MCC gene trees resulting from the constrained gene-tree inference. Outgroup branches are not drawn to scale. H.: Haplochromini; Cyph.: Cyphotilapiini; Lim.: Limnochromini.

The use of concatenated marker sets instead of the multispecies coalescent model led to a number of differences in both the topology and the estimated divergence dates (Supplementary Figure S1; Supplementary File S2). Most notably, *Eretmodus cyanostictus* appeared as the sister to a clade combining the Haplochromini, Ectodini, Limnochromini, and Cyphotilapiini in the species tree based on concatenation (BPP = 1.0), but nested within these tribes in analyses using the multispecies coalescent model. With the exception of outgroup splits, divergences between Lake Tanganyika tribes were always older when using concatenation (Fig. 3). The sequence of divergences between the Haplochromini, Eretmodini, Ectodini, and the combined Limnochromini and Cyphotilapiini appeared far less rapid based on concatenation, with mean age estimates between 10.5 (95% HPD 12.7-9.3 Ma) and 8.7 Ma (95% HPD 10.7-6.9 Ma), as compared to 7.4 (95% HPD 9.1-5.8 Ma) to 7.0 Ma (95% HPD 8.7-5.4 Ma) in the species tree based on the multispecies coalescent model. Similarly, the divergence of Lamprologini and the H-lineage was estimated at 12.2 Ma (95% HPD 14.7-9.7 Ma) based on concatenation, but at 9.2 Ma (95% HPD 11.4-7.1 Ma) when using the multispecies coalescent model. Relative differences between age estimates were greatest for the two young radiations of Lake Malawi and Lake Victoria. The MRCA of the four Lake Malawi species included in our taxon set was estimated at 1.1 Ma (95% HPD 1.5-0.7 Ma) based on concatenation, but around 0.7 Ma (95% HPD 1.2-0.4 Ma) in analyses using the multispecies coalescent model. As our taxon set includes cichlids of both the ‘mbuna’ group and of the genus *Rhamphochromis*, the divergence of these species is likely equivalent to the onset of the Lake Malawi radiation (Joyce et al. 2011; Genner et al. 2015; McGee et al. 2016). The first divergence among the four Lake Victoria species of our taxon set was estimated at 0.9 Ma (95% HPD 1.3-0.6 Ma) in concatenation-based analyses, but only around 191 thousand years ago (ka) (95% HPD 322-40 ka) in the species tree based on the multispecies coalescent model (Fig. 3). With genera *Neochromis, Haplochromis, Labrochromis*, and *Pundamilia* included in our taxon set, this divergence event is also likely to be synonymous with the origin of the superflock endemic to Lake Victoria (Verheyen et al. 2003), excluding older Lake Kivu haplochromines.

**Figure 3:**
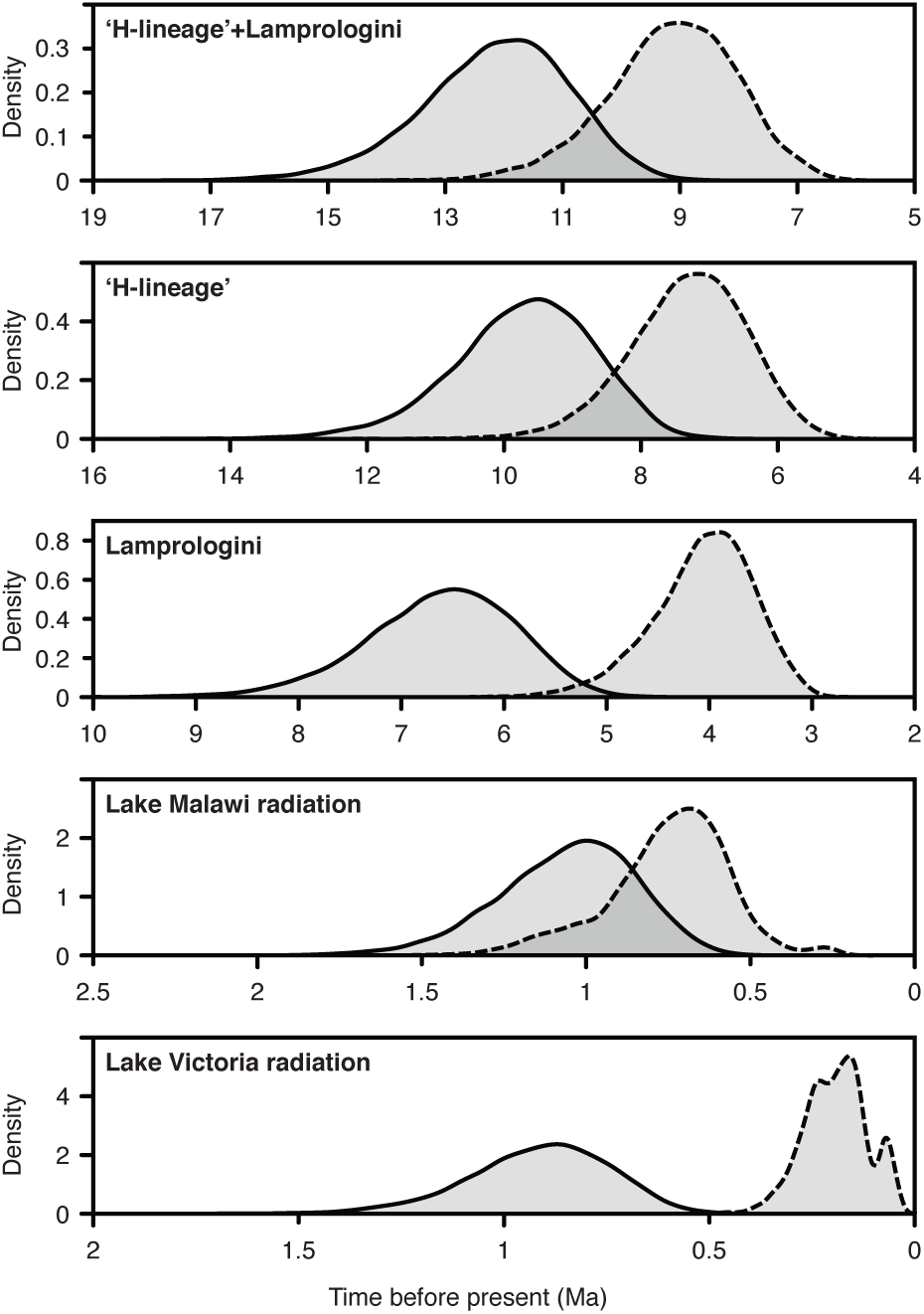
Age estimates for five selected clades of East African cichlids. Posterior age distributions for five selected clades, resulting from Bayesian species-tree analyses based on concatenation (solid black line) or the multispecies coalescent model (dashed black line).

Effective population sizes (*N_e_*) estimated with the multispecies coalescent model ranged between 3.6 x 10^4^ and 8.1 x 10^5^, assuming a mean generation times of three years for cichlid fishes (Malinsky et al. 2015). Notably, two out of the three overall lowest effective population sizes of internal branches were found in the two lineages leading to the Lake Malawi (*N_e_* = 7.5 × 10^4^; 95% HPD 1.4 The timeline of diversification estimated with 10^4^−1.4 x10^5^) and Lake Victoria (*N_e_* = 1.4 × 10^5^; 95% HPD 2.2 × 10^4^−2.7 × 10^5^) radiations. In contrast, high effective population sizes were inferred for the basal lineages of the Lake Tanganyika radiation, with estimates of 6.1 × 10^5^ (95% HPD 2.6 — 9.8 × 10^5^) for the branch leading to the H-lineage, and 3.8 × 10^5^ (95% HPD 2.4 — 5.3 × 10^5^) in the stem lineage of Lamprologini, and 5.1 × 10^5^ (95% HPD 2.9 — 7.3 × 10^5^) in the common ancestor of these two groups.

### Constrained gene-tree inference based on the species tree

Adding species-tree constraints to the inference of individual gene trees greatly improved the species-level node support of these gene trees (mean BPP = 0.64, compared to mean BPP = 0.43 in unconstrained gene-tree analyses). The addition of species putatively involved in introgression to the taxon set resulted in very different attachment points for *Boulengerochromis microlepis, Perissodus microlepis*, and *Pseudocrenilabrus philander* in individual constrained gene trees (Fig. 2; Supplementary File S2). *Boulengerochromis microlepis* was found in a sister-group position to *Trematocara nigrifrons* in ten gene trees, but also appeared more closely related to the Ectodini, the H-lineage, the Lamprologini, or a clade combining the H-lineage and Lamprologini in a total of 18 constrained gene trees. In contrast, *Perissodus microlepis* was most often recovered as the sister of the Ectodini (in ten constrained gene trees), but appeared closely related to the Cyphotilapiini (six constrained gene trees) and Limnochromini (five constrained gene trees), and also diverged from basal branches in several of the constrained gene trees. Attachment points for *Pseudocrenilabrus philander* were mostly found within the Haplochromini (13 constrained gene trees), but the species also appeared as the sister of the Ectodini (six constrained gene trees), the Eretmodini (three constrained gene trees), or the Cyphotilapiini (three constrained gene trees). All attachment points of these three species are shown in Fig. 2.

### Likelihood-based tests for introgression

Maximum likelihood network topologies were recovered consistently in at least nine out of ten analysis replicates for each set of analyses with the software PhyloNet, indicating run convergence. All resulting maximum likelihood network topologies were completely congruent with the previously inferred species tree (Fig. 2), except that *Eretmodus cyanostictus* appeared closer to the Cyphotilapiini and Limnochromini than to the Haplochromini when *Pseudocrenilabrus philander* was included in the taxon set. The addition of single introgression edges to the tree improved the likelihood by 28.5, 14.2, and 2.9 log units when *Boulengerochromis microlepis, Perissodus microlepis*, and *Pseudocrenilabrus philander*, respectively, were included in the taxon set. *Boulengerochromis microlepis* was recovered as the sister lineage of the clade combining the H-lineage and Lamprologini, and appears to have received introgression from the common ancestor of Cyphotilapiini and Limnochromini, with an estimated inheritance probability of 24% (Fig. 4A). *Perissodus microlepis*, on the other hand was found as the sister of Cyphotilapiini and Limnochromini, and a part of its genome appears to have introgressed from the common ancestor of H-lineage and Lamprologini, with a slightly lower inferred inheritance probability of 21% (Fig. 4B). Thus, the two introgression edges inferred for *Boulengerochromis microlepis* and *Perissodus microlepis* are opposing each other, which could be taken as indication that gene flow between the common ancestor of H-lineage and Lamprologini and the common ancestor of Cyphotilapiini and Limnochromini was bidirectional and occurred through hybridization between a member of Boulengerochromini (or a shared ancestor with Bathybatini) and a member of Perissodini (or a shared ancestor with Cyprichromini). Finally, *Pseudocrenilabrus philander* was recovered as the sister lineage of other Haplochromini, and appears to have received introgression from Cyphotilapiini, with an estimated inheritance probability of 9% (Fig. 4C). Thus, we find strong support (≥ 14.2 log-likelihood units) for hypotheses H1 and H2 of introgression between the H-lineage and the Boulengerochromini, and moderate support (2.9 log-likelihood units) for hypothesis H3 of introgression from the Cyphotilapiini to Lake Tanganyika Haplochromini (“Tropheini”).

**Figure 4:**
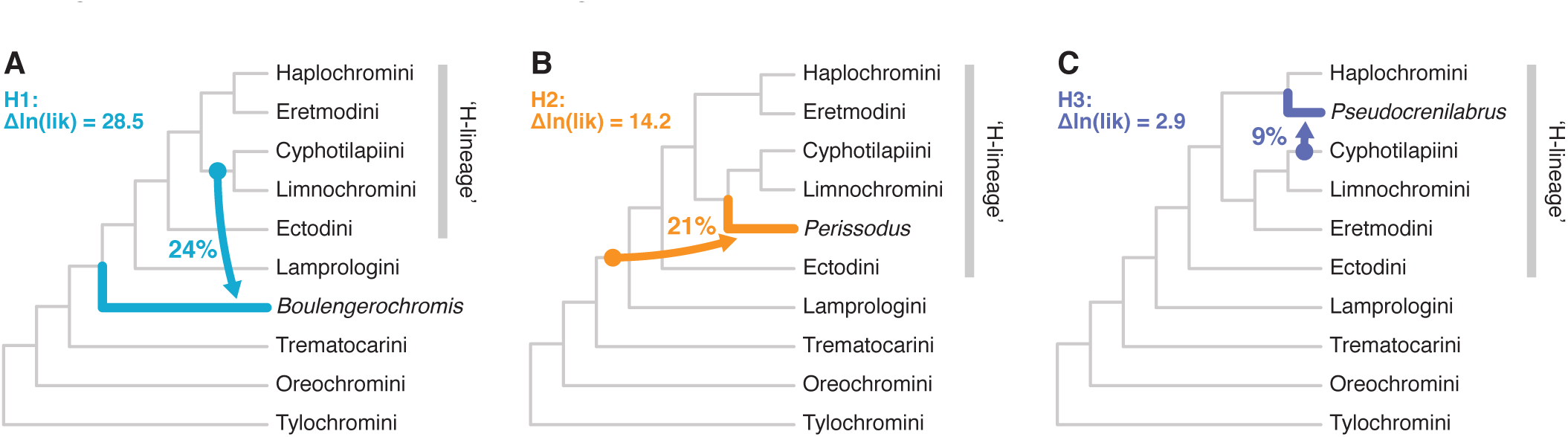
The three inferred introgression events between East Afrcian cichlid tribes. Introgression events inferred with the likelihood framework implemented in PhyloNet. Arrows indicate the direction of introgression events, and percentages given represent inheritance probabilities estimated for introgression edges. The improvement in likelihood gained by addition of this introgression edge is indicated.

## Discussion

### A refined species-tree topology for Lake Tanganyika cichlids

With hundreds of species of fish, mollusks and crustaceans found nowhere else on earth, Lake Tanganyika is a textbook example of the evolution of a complex ecosystem in isolation (Salzburger et al. 2014). In particular, the endemic fauna of cichlid fishes has received considerable scientific attention including phylogenetic estimates using molecular sequence data. Earlier mtDNA-based phylogenetic hypotheses for Lake Tanganyika cichlids — sometimes in combination with a small number of nuclear markers — were characterized by a partial lack of resolution (especially with respect to relationships between tribes) and low support values for many nodes, as well as inconsistencies between studies (Kocher et al. 1995; Salzburger et al. 2002a; Clabaut et al. 2005; Day et al. 2008; Koblmüller et al. 2008; Muschick et al. 2012). The rapid development of sequencing techniques leading to the availability of multiple independent molecular markers, in association with novel algorithms, promises to improve species-tree estimates (Delsuc et al. 2005; Lemmon and Lemmon 2013).

In a recent study (Meyer et al. 2015), we presented a well-supported phylogenetic hypothesis of the relationships between Lake Tanganyika cichlid tribes based on 42 nuclear loci using a concatenation approach. The strategy of concatenating multilocus data has some limitations, though. For example, in situations of prevalent hybridization or incomplete lineage sorting, a concatenation approach might not be able to redraw the correct species history. Incomplete lineage sorting is particularly common in rapidly diverging lineages, such as in adaptive radiations, due to the relatively small number of generations between speciation events (Suh et al. 2015). The degree of incomplete lineage sorting also increases with effective population sizes, which are likely to be high for cichlids in Lake Tanganyika (Koblmüller et al. 2006), and could thus lead to deep coalescence events and gene trees that are anomalous (i.e. incongruent to the actual species tree).

Gene-tree discordance originating from stochastic processes such as incomplete lineage sorting can be accounted for with coalescent-based approaches, which has been demonstrated on theoretical grounds (Kubatko and Degnan 2007; Heled and Drummond 2010), and based on simulated (Leaché and Rannala 2011) and empirical data (Linkem et al. 2016). However, simulation-based studies have also suggested that even coalescent-based species-tree estimates can be misleading when gene flow is present between lineages (Leaché et al. 2014). To overcome this problem, species included in the analysis should be carefully selected to exclude taxa that are affected by introgression. Species-tree estimates can be further improved when multiple individuals per lineage are sampled, as this increases the number of coalescent events used to calculate population sizes (Heled and Drummond 2010; Knowles and Kubatko 2010).

In the present study, we thus increased allele sampling for the taxon set used in Meyer et al. (2015) (we here use up to 16 specimens per species), and applied the multispecies coalescent model to obtain a refined estimation of the species tree of Lake Tanganyika cichlids. To meet the requirements of the multispecies coalescent model, we first determined the set of species with the strongest signals of putative introgression, and excluded these from species-tree analyses. For comparative reasons, we also reconstructed the tree from the same data set but using the concatenation approach.

In the resulting “introgression-free” species tree based on the multispecies coalescent model (Fig. 2), the only representative of the Trematocarini (*Trematocara nigrifrons*) was placed as the sister lineage to a clade formed by the Lamprologini and all H-lineage tribes. Within the H-lineage, Ectodini was recovered as the sister group to a clade formed by Limnochromini and Cyphotilapiini, together with Eretmodini and Haplochromini. The positioning of Eretmodini within the H-lineage — as sister lineage to Haplochromini — is in agreement with previous phylogenetic hypotheses based on allozyme-(Nishida 1991), AFLP-(Weiss et al. 2015), and multilocus data (Friedman et al. 2013; Meyer et al. 2015; McGee et al. 2016), but stands in contrast to inferences based on mtDNA, which placed Eretmodini outside the H-lineage (Kocher et al. 1995; Clabaut et al. 2005; Day et al. 2008; Muschick et al. 2012). In our concatenated analysis, Eretmodini appeared at yet another position, at the base of the H-lineage (Supplementary Figure S1; also see Fig. 2b of Meyer et al. 2015). Based on the observed cytonuclear discordance and the variable placement of Eretmodini, it has previously been suggested that this tribe originated from an ancient hybridization event involving a Lamprologini-like ancestor and a member of the H-lineage (Clabaut et al. 2005; Weiss et al. 2015; Meyer et al. 2015). Our approach of identifying introgression based on MRCA ages in three-taxon comparisons (Fig. 1), *f*-statistics, and likelihood analyses did not retrieve a hybrid signature for *Eretmodus cyanostictus.* At the same time, the placement of Eretmodini as sister lineage to Haplochromini is only relatively weakly supported (Fig. 2), calling for additional analyses using genome-wide data and denser taxon sampling to firmly place this tribe in the phylogeny of East African cichlids.

Overall, however, the application of the multispecies coalescent model, subsequent to the exclusion of the species showing the strongest signals of introgression, resulted in a refined species-tree estimate for Lake Tanganyika cichlids compared to our previous study (Meyer et al. 2015). For example, we recovered strong support for the monophyly of Limnochromini and Cyphotilapiini (see also Friedman et al. 2013; McGee et al. 2016), which both occur in the deep-water habitat of Lake Tanganyika (Coulter 1991). Two other branches remain weakly supported — the one placing Ectodini at the base of the H-lineage and the above-mentioned branch placing Eretmodini as sister lineage to Haplochromini. The apparent lack of phylogenetic resolution among these branches in our analyses with the multispecies coalescent model (Fig. 2), is likely to reflect the rapidity of lineage formation at the base of the H-lineage (see also Salzburger et al. 2002b; Koblmüller et al. 2004; Duftner et al. 2005; Koblmüller et al. 2008). While the basal branches within the H-lineage receive higher support values based on the concatenation model (Supplementary Figure 1), these may be misleading, as concatenation has been shown to overestimate branch support and potentially produce incorrect species-tree estimates (Ogilvie et al. 2016; Linkem et al. 2016). In contrast, using the multispecies coalescent model with only tens of loci may produce more accurate species trees than concatenation with far larger data sets (Ogilvie et al. 2016).

The drawback of our strategy to produce an “introgression-free” species-tree estimate is that it comes on the expense of the exclusion of certain taxa (the putative recipients of introgression). The resulting species tree is therefore robust, yet incomplete. Nevertheless, there are ways to interpret the placement of the taxa that were excluded from species-tree analyses due to signals of introgression. For example, under the assumption that the true phylogenetic signal is stronger than the signal resulting from introgression, the frequencies of the attachment of a lineage to a given branch in the individual gene trees can be considered as democratic vote for its phylogenetic position (indicated by dots on branches in Fig. 2). We exemplify this for the three taxa with the strongest signals of introgression. Based upon the inspection of individual gene trees, *Boulengerochromis microlepis* is suggested to be a sister lineage of *Trematocara nigrifrons* (as e.g. Salzburger et al. 2002b; Clabaut et al. 2005; Meyer et al. 2015), or, alternatively, more closely related to the common ancestor the Lamprologini and the H-lineage (as in Weiss et al. 2015). *Perissodus microlepis* shows affinities to the ancestor of Ectodini (see e.g. Kocher et al. 1995; Clabaut et al. 2005) or is placed in a clade together with Limnochromini and Cyphotilapiini (as observed in Salzburger et al. 2002b; Friedman et al. 2013). Finally, the riverine haplochromine *Pseudocrenilabrus philander* is most often placed near the base of the Haplochromini clade (see e.g. Salzburger et al. 2002b; Clabaut et al. 2005; Santos et al. 2014; Meyer et al. 2015), but also appears frequently as sister lineage to the Ectodini, which has not been observed previously.

### Application of the multispecies coalescent model improves age estimates for East African radiations

The timeline of diversification estimated with the multispecies coalescent model was consistently younger than that obtained with concatenated sequences for all markers. This observation is in agreement with the differences between the models employed by the two approaches. While the concatenation model implicitly assumes that all gene trees are identical to each other as well as to the species tree (Edwards et al. 2016), the multispecies coalescent model accounts for genetic divergence that predates speciation. As a result, concatenation is likely to result in overestimated speciation times, and the degree of this overestimation depends on the ancestral effective population sizes (McCormack et al. 2011). With large population sizes, as are commonly found in adaptive radiations of fishes (Won et al. 2005), the difference in age estimates obtained with the two approaches can be on the order of millions of years (Colombo et al. 2015; Ogilvie et al. 2016). As all previous fossil-based phylogenetic time calibrations of the East African cichlid diversification were based on the concatenation model (e.g. Genner et al. 2007; Schwarzer et al. 2009; Friedman et al. 2013; McMahan et al. 2013), our timeline estimated with the multispecies coalescent model is likely to provide more realistic divergence dates for cichlid radiations in lakes Tanganyika, Malawi, and Victoria.

*Lake Tanganyika radiation*.— Even though geological data suggest that Lake Tanganyika originated around 12-9 Ma (Cohen et al. 1993; Salzburger et al. 2014), concatenation-based age estimates for tribes endemic to this lake are far older. Regardless of whether an assumed Gondwanan vicariance or the cichlid fossil record were used for divergence-time analyses, estimates of Genner et al. (2007) imply that at least 10 cichlid lineages colonized Lake Tanganyika independently but left no traces of their existence outside of the lake. Using calibrations based on the phylogeny of Azuma et al. (2008) and the fossil *Oreochromis \lorenzoi* (Carnevale et al. 2003), Schwarzer et al. (2009) estimated the MRCA of Lamprologini and the H-lineage between 20.4 and 10.6 Ma, with a mean age estimate of 15.4 Ma. This estimate is comparable to our concatenation-based age estimate for the same node (12.2 Ma), but is likely to predate the origin of Lake Tanganyika. In contrast, analyses with the multispecies coalescent model suggest that Lamprologini and the H-lineage diverged around 9.2 Ma, which could thus represent the first divergence event within Lake Tanganyika (Fig. 3). Our timeline is therefore consistent with a colonization history of Lake Tanganyika that requires two to four colonization events from lineages that subsequently went extinct outside the lake (Trematocarini, the common ancestor of Lamprologini and the H-lineage, and possibly the ancestors of Boulengerochromini and Bathybatini) (Salzburger et al. 2002b).

Within the H-lineage, we identified a sequence of rapid splitting events with the multispecies coalescent model that began with the separation of Ectodini around 7.4 Ma and ended with the divergence of Eretmodini and Haplochromini, only about 400 000 years later. In contrast, estimates for these divergence events are stretched out over almost 2 myr when inferred with the concatenation model, which is indicative of a high degree of ILS and thus discordance of individual gene trees. Our coalescent-based age estimates for these lineages coincide with the topographic depression of the northern Lake Tanganyika basin (8-7 Ma; Cohen et al. 1993), suggesting that the main diversification of Lake Tanganyika tribes could have been triggered by ecological opportunity (see e.g. Wagner et al. 2012) in this newly-formed lake basin.

*Lake Malawi radiation*.— Lake Malawi has existed perennially for 4.5-4.0 myr (Ring and Betzler 1995), which is consistent with most phylogenetic age estimages for its largely endemic cichlid radiation (Genner et al. 2007; Schwarzer et al. 2009; Friedman et al. 2013; McMahan et al. 2013; Loh et al. 2013). However, throughout its history, Lake Malawi has experienced severe lake-level fluctuations with lowstands that had considerable impact on its cichlid fauna due to habitat reduction and eutrophication (Salzburger et al. 2014; Lyons et al. 2015). Our age estimate based on the multispecies coalescent model (749 ka) nearly coincides with a shift in hydroclimate regimes during the Mid-Pleistocene Transition (∽ 800 ka) that led to a wetter climate and less frequent lake-level fluctuations (Lyons et al. 2015). These changes could have promoted diversification of Lake Malawi cichlid fishes due to increased ecological opportunity and habitat stability.

*Lake Victoria radiation*.— While the Lake Victoria basin originated around 400 ka (Johnson et al. 1996; Salzburger et al. 2014), the radiation of the endemic Lake Victoria superflock (sensu Verheyen et al. 2003, excluding Lake Kivu species) has long been thought to be younger than 200 ka, based on mitochondrial mutation rate estimates and palaeogeographic considerations (Meyer et al. 1990; Johnson et al. 1996; Verheyen et al. 2003). However, phylogenetic time calibrations have so far failed to reproduce these young age estimates (Elmer et al. 2009; Wagner et al. 2012; Rabosky et al. 2013). Whereas our timeline based on concatenation suggest an early origin of the Lake Victoria superflock around 913 ka, analyses with the multispecies coalescent model support a much younger origin around 191 ka, in agreement with earlier hypotheses (Meyer et al. 1990; Johnson et al. 1996; Verheyen et al. 2003). This indicates that most of the genetic variation present in cichlids of the Lake Victoria superflock pre-existed in the common ancestor. Our estimate of the effective population size of this ancestor (*N_e_* = 143,000) is small compared to those of other internal branches but nevertheless suggests a rather large founding population of the Lake Victoria superflock. This could indicate that Lake Victoria was colonized gradually through previously existing river connections (see Seehausen et al. 2003; Salzburger et al. 2005) rather than by singular dispersal events. It should be noted, however, that effective population sizes estimation with the multispecies coalescent model assumes an absence of population structure and introgression, and that violation of these assumptions may result in overestimated population sizes.

Our phylogentic time calibration was based on secondary constraints taken from the timeline of global cichlid diversification in McMahan et al. (2013) and are thus dependent on the correct estimation of this timeline. According to age estimates of McMahan et al. (2013), Tylochromini and Oreochromini originated around 42 and 23 Ma, respectively, which is in agreement with the earliest fossil records of both tribes, ? *Tylochromis* from the Egyptian Jebel Qatrani Formation (35.12-33.77 Ma; Murray 2002, 2004), and *Oreochromis †martyini* from the Ngorora Formation of Kenya (12.0-9.3 Ma; Van Couvering 1982; Murray and Stewart 1999). Moreover, age estimates of McMahan et al. (2013) were intermediate between those found in other studies, e.g. the age of Tylochromini was estimated between 63.7 and 33.4 Ma in Genner et al. (2007), at around 67 Ma in Azuma et al. (2008), around 56.7 Ma in Schwarzer et al. (2009), and around 24.3 Ma in Friedman et al. (2013). The older estimates of Azuma et al. (2008), Genner et al. (2007), and Schwarzer et al. (2009) all directly or indirectly rely on the assumption of Gondwanan vicariance between Neotropical and African cichlid fishes, which remains debated (Friedman et al. 2013) and disagrees with recent timelines for the diversification of teleost fishes (Near et al. 2012, 2013; Betancur-R et al. 2013; Rabosky et al. 2013). On the other hand, the younger age estimates of Friedman et al. (2013) partially disagree with the fossil record of cichlids, as several clades appear far younger than their first fossil occurrences (Chen et al. 2014; Musilová et al. 2015). Therefore, we suggest that intermediate age estimates based on the timeline of McMahan et al. (2013) may provide the most realistic picture of East African cichlid diversification.

### Introgression in Lake Tanganyika cichlids

The exchange of genetic material between species is a common phenomenon in adaptive radiations (see e.g. Berner and Salzburger 2015), ranging from ongoing gene flow between recently diverged species to more ancient introgression events (The Heliconius Genome Consortium 2012; Jones et al. 2012; Lamichhaney et al. 2015). Likewise, recent and more ancient hybridization events are well documented for East African cichlid fishes (Lake Tanganyika: Koblmüller et al. 2007, 2010; Sturmbauer et al. 2010; Weiss et al. 2015; Lake Malawi: Genner and Turner 2012; Joyce et al. 2011; Lake Victoria: Keller et al. 2013; admixture across watersheds: Schwarzer et al. 2012; Loh et al. 2013). Moreover, it has been suggested that hybridization might promote adaptive radiation in the first place (Grant and Grant 1992; Seehausen 2004) via the creation of novel combinations of parental alleles leading to increased genetic diversity and novel phenotypes in the offspring, upon which natural and sexual selection can act (Hedrick 2013; Arnold 2015). In some situations, e.g. after the colonization of novel habitats, hybrids might be better adapted for the exploitation of new ecological niches (see e.g. Rieseberg et al. 2003; Seehausen 2004; Stelkens and Seehausen 2009; Abbott et al. 2013; Keller et al. 2013; Seehausen et al. 2014).

Here, we applied a novel approach based on MRCA ages in three-taxon comparisons, *f*-statistics, and likelihood analyses to detect introgression between members of different cichlid tribes in Lake Tanganyika. Our analyses uncovered three past introgression events that left behind particularly strong signatures in the genomes of the descendant lineages (Figs. 1,4). The most strongly supported case of past introgression involves the ancestor of *Boulengerochromis microlepis* (or its common ancestor with Bathybatini) as recipient, and an early member of Perissodini or Limnochromini as donor. In a second event, *B. microlepis* (or an earlier member of Boulengerochromini or Bathybatini) served as donor, whereas the common ancestor of Perissodini and Cyprichromini was identified as the most likely recipient. Since both instances involve — in inverted positions — a similar set of recipient and donor lineages, it is possible that they reflect a single introgression event responsible for the reciprocal exchange of substantial portions of genetic material between a more ancient lineage of cichlids (the ancestor of Boulengerochromini or of Boulengerochromini and Bathybatini) and a member of the more derived H-lineage. Thirdly, there is evidence for introgression from an early member of Cyphotilapiini into an ancestor of the Lake Tanganyika haplochromines and the closely related riverine genera *Pseudocrenilabrus* and *Serranochromis* (see also Weiss et al. 2015). Within the clade combining Haplochromini of Lake Tanganyika with the two riverine lineages, the strongest signals of introgression were observed in *Pseudocrenilabrus philander*, closely followed by the Lake Tanganyika endemic *Lobochilotes labiatus.* While we here used the riverine *Pseudocrenilabrus philander* in all analyses of introgression as the representative of this clade (as we consistently chose the taxon with the strongest signals in each case), it seems likely that the introgression event affecting this clade took place within Lake Tanganyika and thus subsequent to its secondary colonization by haplochromines. The presence of introgression signals not only in Lake Tanganyika haplochromines but also in riverine lineages could then be explained by their divergence subsequent to introgression, or by further genetic exchange between these closely related species (Loh et al. 2013). What could have triggered introgressive hybridization between distantly related cichlid lineages (that is, at the level of different tribes) in the confines of Lake Tanganyika, and under which circumstances and environmental conditions such rare hybridization events might have taken place, remains a matter of speculation. There are, however, arguments that render past gene exchange between *B. microlepis* and a member of the H-lineage, and between Cyphotilapiini and Haplochromini, not entirely improbable. *Boulengerochromis microlepis*, the only member of Boulengerochromini, is unusual in various aspects. Originally assigned to Tilapiini (Poll 1986), *B. microlepis* was put into its own tribe because of its distinctiveness with respect to any other cichlid species known (Takahashi 2003). *Boulengerochromis microlepis* is the largest cichlid in the world, reaching an adult size of ∽80 cm, and it forms breeding pairs for unusually long periods of time, until the offspring has almost reached adult size (Konings 2015). Our results suggest that *B. microlepis* is characterized by a combination of genetic material from a pre-Tanganyikan cichlid lineage and a member of the endemic H-lineage. Thus, gene exchange with lake-adapted lineages might have contributed to the survival of the ancestor of *B. microlepis* after its colonization of this novel environment.

The Tanganyikan representatives of the Haplochromini, the “Tropheini”, are also somewhat unusual compared to other members of that tribe. For example, most “Tropheini” species are much less sexually dimorphic than e.g. the haplochromines of the adaptive radiations in lakes Victoria and Malawi (Fryer and Iles 1972; Salzburger et al. 2006), and some of the Tanganyikan haplochromines become unusually large (up to 40 cm in length and 3 kg in size the case of *Petrochromis* sp. ‘giant’). Interestingly, there are morphological similarities between the Tanganyikan haplochromines and the Cyphotilapiini. In fact, both genera currently included in Cyphotilapiini, *Cyphotilapia* and *Trematochromis*, have initially been grouped into the Haplochromini based on morphological grounds (Poll 1986), and were later suggested to form their own, respectively monotypic tribes because of their phenotypic distinctiveness (Takahashi 2003). The only representative of the latter genus (*Trematochromis benthicola*) was, originally, even placed into the same genus as the Tanganyikan haplochromine *Ctenochromis horei* and has only recently been established as a member of the Cyphotilapiini based on molecular data (Muschick et al. 2012).

In the light of our new results suggesting ancient gene flow from Cyphotilapiini into Haplochromini, the morphological resemblance between the members of these tribes can be re-interpreted as being due to past introgression (see e.g. The Heliconius Genome Consortium 2012). Our data place the introgression event from Cyphotilapiini to Haplochromini near the common ancestor of the “Tropheini” and riverine lineages (*Pseudocrenilabrus* and *Serranochromis*), while other, more derived, Haplochromini of Lake Malawi and Lake Victoria do not seem to be affected. This suggests that the radiation of the haplochromines in Lake Tanganyika started out from a riverine ancestor that invaded the lake secondarily and hybridized with a lake-adapted species. This scenario is in line with the idea that hybridization can trigger adaptive radiation (Seehausen 2004), e.g. through transgressive segregation (Stelkens and Seehausen 2009). Unlike in the haplochromine adaptive radiations in lakes Victoria and Malawi, for which precursory hybridization between exclusively riverine lineages has been proposed (Seehausen et al. 2003; Joyce et al. 2011), the Tanganyikan haplochromines appear to be the product of a riverine and a lake-adapted form.

The here reported cases of past introgression between different cichlid tribes in Lake Tanganyika have in common an asymmetric fate with respect to taxonomic diversity of the two lineages involved in the exchange of genetic material. Each event brought forth an ancestor of a lineage that subsequently diversified into an array of species (Perissodini and Cyprichromini: 16 species; Haplochromini: ∽30 species in Lake Tanganyika), while leaving behind another lineage with low species diversity (Cyphotilapiini: 3 species; Boulengerochromini: 1 species). What might have caused this asymmetry in speciation rates after gene exchange, and how introgression is connected to diversification overall in Lake Tanganyika cichlids, should be in the focus of future studies. In this context, it is interesting to note that we did not find evidence for inter-tribal introgression events that involve a member of the most species-rich cichlid tribe in Lake Tanganyika, Lamprologini, although — within this tribe — introgression and hybridization appear to be common (Salzburger et al. 2002a; Sturmbauer et al. 2010). A possible explanation for the “genomic demarcation” of lamprologines from other cichlid lineages in Lake Tanganyika is their distinctive breeding mode: Lamprologines are substrate spawners with only very subtle, if any, phenotypic differences between males and females, whereas all but one (*Boulengerochromis microlepis*) of the remaining Tanganyikan cichlid species are mouth-brooders featuring, in many cases, a pronounced sexual (color) dimorphism (Konings 2015).

Taken together, we uncovered several instances of past inter-tribal introgression in Lake Tanganyika. These involve ancestors of tribes that, until today, could not have been placed firmly into the phylogeny of East African cichlids, suggesting that their chimeric nature is responsible for the difficulties in resolving their respective placements in previous analyses based on both molecular and morphological data. Our identification of multiple introgression events in Lake Tanganyika cichlids is in line with expectations regarding the occurrence of hybridization in adaptive radiation and corroborates the importance of between-species genetic exchange for the evolution of biodiversity.

## Supplementary Material

Sequence data has been deposited on GenBank (www.ncbi.nlm.nih.gov/Genbank) under accession numbers XXXXXX-XXXXXX, and alignment files in Nexus format as well as MCC trees are available from the Dryad Digital Repository: http://dx.doi.org/10.5061/dryad.fNNNN].

## Acknowledgements

We would like to thank Marco Colombo, Bernd Egger, Adrian Indermaur, Moritz Muschick, and Frauke Münzel, Fabrizia Ronco, and Anya Theis for assistance during fieldwork, and Ole Seehausen for providing tissue samples; Brigitte Aeschbach and Nicolas Boileau for help with labwork; Christof Wunderlin and Georges Wigger from Microsynth for 454 sequencing; and Alexei Drummond, Thomas Marcussen, Caleb McMahan, and Huw Ogilvie for discussion.

## References

Abbott, R., D. Albach, S. Ansell, J. W. Arntzen, S. J. E. Baird, N. Bierne, J. Boughman, A. Brelsford, C. A. Buerkle, R. Buggs, R. K. Butlin, U. Dieckmann, F. Eroukhmanoff, A. Grill, S. H. Cahan, J. S. Hermansen, G. Hewitt, A. G. Hudson, C. D. Jiggins, J. Jones, B. Keller, T. Marczewski, J. Mallet, P. Martinez-Rodriguez, M. Möst, S. Mullen, R. Nichols, A. W. Nolte, C. Parisod, K. Pfennig, A. M. Rice, M. G. Ritchie, B. Seifert, C. M. Smadja, R. Stelkens, J. M. Szymura, R. Väinölä, J. B. W. Wolf, and D. Zinner. 2013. Hybridization and speciation. J. Evol. Biol. 26:229–246.

Arnold, M. L. 2015. Divergence with Genetic Exchange. Oxford University Press, Oxford, UK.

Azuma, Y., Y. Kumazawa, M. Miya, K. Mabuchi, and M. Nishida. 2008. Mitogenomic evaluation of the historical biogeography of cichlids toward reliable dating of teleostean divergences. BMC Evol. Biol. 8:215.

Bell, M. A. and S. A. Foster. 1994. The Evolutionary Biology of the Threespine Stickleback. Oxford University Press, Oxford, UK.

Berner, D. and W. Salzburger. 2015. The genomics of organismal diversification illuminated by adaptive radiations. Trends Genet. 31:491–499.

Betancur-R, R., R. E. Broughton, E. O. Wiley, K. E. Carpenter, J. A. López, C. Li, N. I. Holcroft, D. Arcila, M. D. Sanciangco, J. C. Cureton, F. Zhang, T. Buser, M. A. Campbell, J. A. Ballesteros, A. Roa-Varon, S. C. Willis, W. C. Borden, T. Rowley, P. C. Reneau, D. J. Hough, G. Lu, T. Grande, G. Arratia, and G. Ortí. 2013. The Tree of Life and a new classification of bony fishes. PLOS Currents: Tree of Life Pages 1–45.

Bouckaert, R., J. Heled, D. Kühnert, T. Vaughan, C.-H. Wu, D. Xie, M. A. Suchard, A. Rambaut, and A. J. Drummond. 2014. BEAST 2: a software platform for Bayesian evolutionary analysis. PLOS Comput. Biol. 10:e1003537.

Carnevale, G., C. Sorbini, and W. Landini. 2003. Oreochromis lorenzoi, a new species of tilapiine cichlid from the Late Miocene of central Italy. J. Vert. Paleontol. 23:508–516.

Chen, W.-J., F. Santini, G. Carnevale, J.-N. Chen, S.-H. Liu, S. Lavoué, and R. L. Mayden. 2014. New insights on early evolution of spiny-rayed fishes (Teleostei: Acanthomorpha). Front. Mar. Sci. 1:53.

Clabaut, C., W. Salzburger, and A. Meyer. 2005. Comparative phylogenetic analyses of the adaptive radiation of Lake Tanganyika cichlid fish: nuclear sequences are less homoplasious but also less informative than mitochondrial DNA. J. Mol. Evol. 61:666–681.

Cohen, A. S., M. J. Soreghan, and C. A. Scholz. 1993. Estimating the age of formation of lakes: An example from Lake Tanganyika, East African Rift system. Geology 21:511–514.

Colombo, M., M. Damerau, R. Hanel, W. Salzburger, and M. Matschiner. 2015. Diversity and disparity through time in the adaptive radiation of Antarctic notothenioid fishes. J. Evol. Biol. 28:376–394.

Coulter, G. W. 1991. The benthic fish community. Pages 151–199 in Lake Tanganyika and its Life (G. W. Coulter, ed.). Oxford University Press, London.

Coyne, J. A. and H. A. Orr. 2004. Speciation. Sinauer Associates, Inc., Sunderland, Massachusetts, USA.

Day, J. J., J. A. Cotton, and T. G. Barraclough. 2008. Tempo and mode of diversification of Lake Tanganyika cichlid fishes. PLOS ONE 3:e1730.

Delsuc, F., H. Brinkmann, and H. Philippe. 2005. Phylogenomics and the reconstruction of the tree of life. Nat. Rev. Genet. 6:361.

Duftner, N., S. Koblmüller, and C. Sturmbauer. 2005. Evolutionary relationships of the Limnochromini, a tribe of benthic deepwater cichlid fish endemic to Lake Tanganyika, East Africa. J. Mol. Evol. 60:277–289.

Dunz, A. R. and U. K. Schliewen. 2013. Molecular phylogeny and revised classification of the haplotilapiine cichlid fishes formerly referred to as “*Tilapia*”. Mol. Phylogenet. Evol. 68:64–80.

Edwards, S. V., Z. Xi, A. Janke, B. C. Faircloth, J. E. McCormack, T. C. Glenn, B. Zhong, S. Wu, E. M. Lemmon, A. R. Lemmon, A. D. Leaché, L. Liu, and C. C. Davis. 2016. Implementing and testing the multispecies coalescent model: A valuable paradigm for phylogenomics. Mol. Phylogenet. Evol. 94:447–462.

Elmer, K. R., C. Reggio, T. Wirth, E. Verheyen, W. Salzburger, and A. Meyer. 2009. Pleistocene desiccation in East Africa bottlenecked but did not extirpate the adaptive radiation of Lake Victoria haplochromine cichlid fishes. Proc. Natl. Acad. Sci. USA 106:13404–13409.

Excoffier, L., I. Dupanloup, E. Huerta-Sañchez, V. C. Sousa, and M. Foll. 2013. Robust demographic inference from genomic and SNP data. PLOS Genet. 9:e1003905.

Friedman, M., B. P. Keck, A. Dornburg, R. I. Eytan, C. H. Martin, C. D. Hulsey, P. C. Wainwright, and T. J. Near. 2013. Molecular and fossil evidence place the origin of cichlid fishes long after Gondwanan rifting. Proc. R. Soc. B 280:20131733.

Fryer, G. and T. D. Iles. 1972. The Cichlid Fishes of the Great Lakes of Africa: Their Biology and Evolution. Oliver and Boyd Press, Scotland.

Gavryushkina, A., D. Welch, T. Stadler, and A. J. Drummond. 2014. Bayesian inference of sampled ancestor trees for epidemiology and fossil calibration. PLOS Comput. Biol. 10:e1003919.

Genner, M. J., B. P. Ngatunga, S. Mzighani, A. Smith, and G. F. Turner. 2015. Geographical ancestry of Lake Malawi’s cichlid fish diversity. Biology Letters 11:20150232.

Genner, M. J., O. Seehausen, D. H. Lunt, D. A. Joyce, P. W. Shaw, G. R. Carvalho, and G. F. Turner. 2007. Age of cichlids: new dates for ancient lake fish radiations. Mol. Biol. Evol. 24:1269–1282.

Genner, M. J. and G. F. Turner. 2012. Ancient hybridization and phenotypic novelty within Lake Malawi’s cichlid fish radiation. Mol. Biol. Evol. 29:195–206.

Glor, R. E. 2010. Phylogenetic insights on adaptive radiation. Annu. Rev. Ecol. Evol. Syst. 41:251–270.

Grant, P. and B. Grant. 2008. How and why species multiply: the radiation of Darwin’s finches. Princeton University Press Princeton, New Jersey.

Grant, P. R. and B. R. Grant. 1992. Hybridization of bird species. Science 256:193–197.

Hasegawa, M., H. Kishino, and T. Yano. 1985. Dating of the human-ape splitting by a molecular clock of mitochondrial DNA. J. Mol. Evol. 22:160–174.

Hedrick, P. W. 2013. Adaptive introgression in animals: examples and comparison to new mutation and standing variation as sources of adaptive variation. Mol. Ecol. 22:4606–4618.

Heled, J. and A. J. Drummond. 2010. Bayesian inference of species trees from multilocus data. Mol. Biol. Evol. 27:570–580.

Johnson, T., C. Scholz, M. Talbot, K. Kelts, R. Ricketts, G. Ngobi, K. Beuning, I Ssemmanda, and J. McGill. 1996. Late Pleistocene desiccation of Lake Victoria and rapid evolution of cichlid fishes. Science 273:1091–1093.

Jones, F. C., M. G. Grabherr, Y. F. Chan, P. Russell, E. Mauceli, J. Johnson, R. Swofford, M. Pirun, M. C. Zody, S. White, E. Birney, S. Searle, J. Schmutz, J. Grimwood, M. C. Dickson, R. M. Myers, C. T. Miller, B. R. Summers, A. K. Knecht, S. D. Brady, H. Zhang, A. A. Pollen, T. Howes, C. Amemiya, J. Baldwin, T. Bloom, D. B. Jaffe, R. Nicol, J. Wilkinson, E. S. Lander, F. Di Palma, K. Lindblad-Toh, and D. M. Kingsley. 2012. The genomic basis of adaptive evolution in threespine sticklebacks. Nature 484:55–61.

Joyce, D. A., D. H. Lunt, M. J. Genner, G. F. Turner, R Bills, and O. Seehausen. 2011. Repeated colonization and hybridization in Lake Malawi cichlids. Curr. Biol. 21:R108–R109.

Katoh, K. and D. M. Standley. 2013. MAFFT multiple sequence alignment software version 7: improvements in performance and usability. Mol. Biol. Evol. 30:772–780.

Keller, I., C. E. Wagner, L. Greuter, S. Mwaiko, O. M. Selz, A. Sivasundar, S. Wittwer, and O. Seehausen. 2013. Population genomic signatures of divergent adaptation, gene flow and hybrid speciation in the rapid radiation of Lake Victoria cichlid fishes. Mol. Ecol. 22:2848–2863.

Knowles, L. L. and L. S. Kubatko, eds. 2010. Estimating Species Trees: Practical and Theoretical Aspects. Wiley-Blackwell, Hoboken, NJ, USA.

Koblmüller, S., N. Duftner, K. M. Sefc, M. Aibara, M. Stipacek, M. Blanc, B Egger, and C. Sturmbauer. 2007. Reticulate phylogeny of gastropod-shell-breeding cichlids from Lake Tanganyika - the result of repeated introgressive hybridization. BMC Evol. Biol. 7:7.

Koblmüller, S., B. Egger, C Sturmbauer, and K. M. Sefc. 2010. Rapid radiation, ancient incomplete lineage sorting and ancient hybridization in the endemic Lake Tanganyika cichlid tribe Tropheini. Mol. Phylogenet. Evol. 55:318–334.

Koblmüller, S., W. Salzburger, and C. Sturmbauer. 2004. Evolutionary relationships in the sand-dwelling cichlid lineage of Lake Tanganyika suggest multiple colonization of rocky habitats and convergent origin of biparental mouthbrooding. J. Mol. Evol. 58:79–96.

Koblmüller, S., U. K. Schliewen, N. Duftner, K. M. Sefc, C Katongo, and C. Sturmbauer. 2008. Age and spread of the haplochromine cichlid fishes in Africa. Mol. Phylogenet. Evol. 49:153–169.

Koblmüller, S., K. M. Sefc, N. Duftner, M Warum, and C. Sturmbauer. 2006. Genetic population structure as indirect measure of dispersal ability in a Lake Tanganyika cichlid. Genetica 130:121–131.

Koch, M., S. Koblmüller, K. M. Sefc, N. Duftner, C Katongo, and C. Sturmbauer. 2007. Evolutionary history of the endemic Lake Tanganyika cichlid fish *Tylochromis polylepis*: a recent intruder to a mature adaptive radiation. J. Zool. Syst. Evol. Res. 45:64–71.

Kocher, T. D. 2004. Adaptive evolution and explosive speciation: the cichlid fish model. Nat. Rev. Genet. 5:288–298.

Kocher, T. D., J. A. Conroy, K. R. McKaye, J. R. Stauffer, and S. F. Lockwood. 1995. Evolution of NADH dehydrogenase subunit 2 in East African cichlid fish. Mol. Phylogenet. Evol. 4:420–432.

Konings, A. 2015. Tanganyika cichlids in their natural habitat (3rd edition). Cichlid Press, El Paso, Texas, USA.

Kubatko, L. S. and J. H. Degnan. 2007. Inconsistency of phylogenetic estimates from concatenated data under coalescence. Syst. Biol. 56:17–24.

Lamichhaney, S., J. Berglund, M. S. Almén, K. Maqbool, M. Grabherr, A. Martinez-Barrio, M. Promerová, C.-J. Rubin, C. Wang, N. Zamani, B. R. Grant, P. R. Grant, M. T. Webster, and L. Andersson. 2015. Evolution of Darwin’s finches and their beaks revealed by genome sequencing. Nature 518:371–375.

Leaché, A. D., R. B. Harris, B Rannala, and Z. Yang. 2014. The influence of gene flow on species tree estimation: a simulation study. Syst. Biol. 63:17–30.

Leaché, A. D. and B. Rannala. 2011. The accuracy of species tree estimation under simulation: a comparison of methods. Syst. Biol. 60:126–137.

Lemmon, E. M. and A. R. Lemmon. 2013. High-throughput genomic data in systematics and phylogenetics. Annu. Rev. Ecol. 44:99–121.

Li, H. and R. Durbin. 2010. Fast and accurate long-read alignment with Burrows-Wheeler transform. Bioinformatics 26:589–595.

Linkem, C. W., V. N. Minin, and A. D. Leaché. 2016. Detecting the anomaly zone in species trees and evidence for a misleading signal in higher-level skink phylogeny (Squamata: Scincidae). Syst. Biol. Advance Access, doi:10.1093/sysbio/syw001.

Loh, Y. H. E., E. Bezault, F. M. Muenzel, R. B. Roberts, R. Swofford, M. Barluenga, C. E. Kidd, A. E. Howe, F. Di Palma, K. Lindblad-Toh, J. Hey, O. Seehausen, W. Salzburger, T. D. Kocher, and J. T. Streelman. 2013. Origins of shared genetic variation in African cichlids. Mol. Biol. Evol. 30:906–917.

Losos, J. 2009. Lizards in an Evolutionary Tree. University of California Press Berkeley, California, USA.

Lyons, R. P., C. A. Scholz, A. S. Cohen, J. W. King, E. T. Brown, S. J. Ivory, T. C. Johnson, A. L. Deino, P. N. Reinthal, M. M. McGlue, and M. W. Blome. 2015. Continuous 1.3-million-year record of East African hydroclimate, and implications for patterns of evolution and biodiversity. Proc. Natl. Acad. Sci. USA 112:15568–15573.

Malinsky, M., R. J. Challis, A. M. Tyers, S. Schiffels, Y. Terai, B. P. Ngatunga, E. A. Miska, R. Durbin, M. J. Genner, and G. F. Turner. 2015. Genomic islands of speciation separate cichlid ecomorphs in an East African crater lake. Science 350:1493–1498.

Marcussen, T., S. R. Sandve, L. Heier, M. Spannagl, M. Pfeifer, International Wheat Genome Sequencing Consortium, K. S. Jakobsen, B. B. H. Wulff, B. Steuernagel, K. F. X. Mayer, and O.-A. Olsen. 2014. Ancient hybridizations among the ancestral genomes of bread wheat. Science 345:1250092.

Martin, C. H., J. S. Cutler, J. P. Friel, C. Dening Touokong, G. Coop, and P. C. Wainwright. 2015. Complex histories of repeated gene flow in Cameroon crater lake cichlids cast doubt on one of the clearest examples of sympatric speciation. Evolution 69:1406–1422.

Mayr, E. 2001. What evolution is. Basic Books, New York, NY, USA.

McCormack, J. E., J. Heled, K. S. Delaney, A. T. Peterson, and L. L. Knowles. 2011. Calibrating divergence times on species trees versus gene trees: implications for speciation history of *Aphelocoma* jays. Evolution 65:184–202.

McCune, A. R. and N. R. Lovejoy. 1998. The relative rate of sympatric and allopatric speciation in fishes: Tests using DNA sequence divergence between sister species and among clades. Pages 172–185 in Endless Forms: Species and Speciation (D. Howard and S. Berloccher, eds.). Oxford University Press, Oxford, UK.

McGee, M. D., B. C. Faircloth, S. R. Borstein, J. Zheng, C. Darrin Hulsey, P. C. Wainwright, and M. E. Alfaro. 2016. Replicated divergence in cichlid radiations mirrors a major vertebrate innovation. Proc. R. Soc. B 283:20151413.

McMahan, C. D., P. Chakrabarty, J. S. Sparks, W. L. Smith, and M. P. Davis. 2013. Temporal patterns of diversification across global cichlid biodiversity (Acanthomorpha: Cichlidae). PLOS ONE 8:e71162.

Meyer, A., T. D. Kocher, P. Basasibwaki, and A. C. Wilson. 1990. Monophyletic origin of Lake Victoria cichlid fishes suggested by mitochondrial DNA sequences. Nature 347:550–553.

Meyer, B. S., M. Matschiner, and W. Salzburger. 2015. A tribal level phylogeny of Lake Tanganyika cichlid fishes based on a genomic multi-marker approach. Mol. Phylogenet. Evol. 83C:56–71.

Meyer, B. S. and W. Salzburger. 2012. A novel primer set for multilocus phylogenetic inference in East African cichlid fishes. Mol. Ecol. Resour. 12:1097–1104.

Murray, A. M. 2002. Lower pharyngeal jaw of a cichlid fish (Actinopterygii; Labroidei) from an Early Oligocene site in the Fayum, Egypt. J. Vert. Paleontol. 22:453–455.

Murray, A. M. 2004. Late Eocene and early Oligocene teleost and associated ichthyofauna of the Jebel Qatrani Formation, Fayum, Egypt. Palaeontology 47:711–724.

Murray, A. M. and K. M. Stewart. 1999. A new species of tilapiine cichlid from the Pliocene, Middle Awash, Ethiopia. J. Vert. Paleontol. 19:293–301.

Muschick, M., A. Indermaur, and W. Salzburger. 2012. Convergent evolution within an adaptive radiation of cichlid fishes. Curr. Biol. 22:2362–2368.

Musilová, Z., O. Říčan, Š. Říčanová, P. Janšta, and O. G. J. Novák. 2015. Phylogeny and historical biogeography of trans-Andean cichlid fishes (Teleostei: Cichlidae). Vertebr. Zool. 65:333–350.

Near, T. J., A. Dornburg, R. I. Eytan, B. P. Keck, W. L. Smith, K. L. Kuhn, J. A. Moore, S. A. Price, F. T. Burbrink, M. Friedman, and P. C. Wainwright. 2013. Phylogeny and tempo of diversification in the superradiation of spiny-rayed fishes. Proc. Natl. Acad. Sci. USA 110:12738–12743.

Near, T. J., R. I. Eytan, A. Dornburg, K. L. Kuhn, J. A. Moore, M. P. Davis, P. C. Wainwright, M Friedman, and W. L. Smith. 2012. Resolution of ray-finned fish phylogeny and timing of diversification. Proc. Natl. Acad. Sci. USA 109:13698–13703.

Nishida, M. 1991. Lake Tanganyika as an evolutionary reservoir of old lineages of East African cichlid fishes: inferences from allozyme data. Experientia 47:974–979.

Ogilvie, H. A., J. Heled, D. Xie, and A. J. Drummond. 2016. Computational performance and statistical accuracy of *BEAST and comparisons with other methods. Syst. Biol. Advance Access, doi:10.1093/sysbio/syv118:1-52.

Poll, M. 1986. Classification des Cichlidae du lac Tanganika. Tribus, genres et especes. Académie Royale de Belgique. Classe des Sciences. Mémoires 45:1–163.

Rabosky, D. L., F. Santini, J. Eastman, S. A. Smith, B. Sidlauskas, J. Chang, and M. E. Alfaro. 2013. Rates of speciation and morphological evolution are correlated across the largest vertebrate radiation. Nat. Commun. 4:1958.

Reich, D., K. Thangaraj, N. Patterson, A. L. Price, and L. Singh. 2009. Reconstructing Indian population history. Nature 461:489–494.

Rieseberg, L. H., O. Raymond, D. M. Rosenthal, Z. Lai, K. Livingstone, T. Nakazato, J. L. Durphy, A. E. Schwarzbach, L. A. Donovan, and C. Lexer. 2003. Major ecological transitions in wild sunflowers facilitated by hybridization. Science 301:1211–1216.

Ring, U. and C. Betzler. 1995. Geology of the Malawi Rift: kinematic and tectonosedimentary background to the Chiwondo Beds, northern Malawi. J. Hum. Evol. 28:7–21.

Rokas, A. and S. B. Carroll. 2006. Bushes in the Tree of Life. PLOS Biol. 4:e352.

Salzburger, W. 2009. The interaction of sexually and naturally selected traits in the adaptive radiations of cichlid fishes. Mol. Ecol. 18:169–185.

Salzburger, W., S. Baric, and C. Sturmbauer. 2002a. Speciation via introgressive hybridization in East African cichlids? Mol. Ecol. 11:619–625.

Salzburger, W., T. Mack, E. Verheyen, and A. Meyer. 2005. Out of Tanganyika: Genesis, explosive speciation, key-innovations and phylogeography of the haplochromine cichlid fishes. BMC Evol. Biol. 5:17.

Salzburger, W., A. Meyer, S. Baric, E. Verheyen, and C. Sturmbauer. 2002b. Phylogeny of the Lake Tanganyika cichlid species flock and its relationship to the central and East African haplochromine cichlid fish faunas. Syst. Biol. 51:113–135.

Salzburger, W., H. Niederstatter, A. Brandstatter, B. Berger, W. Parson, J Snoeks, and C. Sturmbauer. 2006. Colour-assortative mating among populations of *Tropheus moorii*, a cichlid fish from Lake Tanganyika, East Africa. Proc. R. Soc. B 273:257–266.

Salzburger, W., B. Van Bocxlaer, and A. S. Cohen. 2014. Ecology and evolution of the African Great Lakes and their faunas. Annu. Rev. Ecol. Evol. Syst. 45:519–545.

Santos, M. E., I. Braasch, N. Boileau, B. S. Meyer, L. Sauteur, A. Böhne, H.-G. Belting, M Affolter, and W. Salzburger. 2014. The evolution of cichlid fish egg-spots is linked with a cis-regulatory change. Nat. Commun. 5:1–11.

Schluter, D. 2000. The Ecology of Adaptive Radiation. Oxford University Press, New York, NY, USA.

Schmieder, R. and R. Edwards. 2011. Quality control and preprocessing of metagenomic datasets. Bioinformatics 27:863–864.

Schwarzer, J., B. Misof, D. Tautz, and U. K. Schliewen. 2009. The root of the East African cichlid radiations. BMC Evol. Biol. 9:186.

Schwarzer, J., E. R. Swartz, E. Vreven, J. Snoeks, F. P. D. Cotterill, B. Y. Misof, and U. K. Schliewen. 2012. Repeated trans-watershed hybridization among haplochromine cichlids (Cichlidae) was triggered by Neogene landscape evolution. Proc. R. Soc. B 279:4389–4398.

Seehausen, O. 2004. Hybridization and adaptive radiation. Trends Ecol. Evol. 19:198–207.

Seehausen, O. 2006. African cichlid fish: a model system in adaptive radiation research. Proc. R. Soc. B 273:1987–1998.

Seehausen, O., R. K. Butlin, I. Keller, C. E. Wagner, J. W. Boughman, P. A. Hohenlohe, C. L. Peichel, G.-P. Saetre, C Bank, Å. Brännström, A. Brelsford, C. S. Clarkson, F. Eroukhmanoff, J. L. Feder, M. C. Fischer, A. D. Foote, P. Franchini, C. D. Jiggins, F. C. Jones, A. K. Lindholm, K. Lucek, M. E. Maan, D. A. Marques, S. H. Martin, B. Matthews, J. I. Meier, M. Möst, M. W. Nachman, E. Nonaka, D. J. Rennison, J. Schwarzer, E. T. Watson, A. M. Westram, and A. Widmer. 2014. Genomics and the origin of species. Nat. Rev. Genet. 15:176–192.

Seehausen, O., E. Koetsier, M. V. Schneider, L. J. Chapman, C. A. Chapman, M. E. Knight, G. F. Turner, J. J. M. v. Alphen, and R. Bills. 2003. Nuclear markers reveal unexpected genetic variation and a Congolese-Nilotic origin of the Lake Victoria cichlid species flock. Proc. R. Soc. B 270:129–137.

Simpson, G. G. 1953. The Major Features of Evolution. Columbia University Press, New York, USA.

Stelkens, R. B. and O. Seehausen. 2009. Genetic distance between species predicts novel trait expression in their hybrids. Evolution 63:884–897.

Sturmbauer, C. and A. Meyer. 1993. Mitochondrial phylogeny of the endemic mouthbrooding lineages of cichlid fishes from Lake Tanganyika in eastern Africa. Mol. Biol. Evol. 10:751–768.

Sturmbauer, C., W. Salzburger, N. Duftner, R Schelly, and S. Koblmüller. 2010. Evolutionary history of the Lake Tanganyika cichlid tribe Lamprologini (Teleostei: Perciformes) derived from mitochondrial and nuclear DNA data. Mol. Phylogenet. Evol. 57:266–284.

Suh, A., L. Smeds, and H. Ellegren. 2015. The dynamics of incomplete lineage sorting across the ancient adaptive radiation of Neoavian birds. PLOS Biol. 13:e1002224.

Takahashi, T. 2003. Systematics of Tanganyikan cichlid fishes (Teleostei: Perciformes). Ichthyol. Res. 50:367–382.

Than, C., D. Ruths, and L. Nakhleh. 2008. PhyloNet: a software package for analyzing and reconstructing reticulate evolutionary relationships. BMC Bioinformatics 9:322.

The Heliconius Genome Consortium. 2012. Butterfly genome reveals promiscuous exchange of mimicry adaptations among species. Nature 487:94–98.

Van Couvering, J. A. H. 1982. Fossil cichlid fish of Africa. Special Papers in Paleontology, The Paleontological Association 29:103.

Verheyen, E., W. Salzburger, J Snoeks, and A. Meyer. 2003. Origin of the superflock of cichlid fishes from Lake Victoria, East Africa. Science 300:325–329.

Wagner, C. E., L. J. Harmon, and O. Seehausen. 2012. Ecological opportunity and sexual selection together predict adaptive radiation. Nature 487:366–369.

Weiss, J. D., F. P. D. Cotterill, and U. K. Schliewen. 2015. Lake Tanganyika—A ‘Melting Pot’ of ancient and young cichlid lineages (Teleostei: Cichlidae)? PLOS ONE 10:e0125043.

Won, Y.-J., A. Sivasundar, Y. Wang, and J. Hey. 2005. On the origin of Lake Malawi cichlid species: A population genetic analysis of divergence. Proc. Natl. Acad. Sci. USA 102:6581–6586.

Yu, Y., J. Dong, K. J. Liu, and L. Nakhleh. 2014. Maximum likelihood inference of reticulate evolutionary histories. Proc. Natl. Acad. Sci. USA 111:16448–16453.

Yule, G. U. 1925. A mathematical theory of evolution, based on the conclusions of Dr. J. C. Willis, F.R.S. Phil. Trans. R. Soc. B 213:21–87.

